# MEK-dependent bioenergetic demand drives terminal CD8^+^ T cell exhaustion

**DOI:** 10.1101/2025.04.30.651453

**Authors:** Tanmana Mitra, Jahan Rahman, Madeline Hwee, Ruben Jose Jesus Faustino Ramos, Hui Liu, Travis Hartman, Justin Cross, Miguel de Jesus, Morgan Huse, Valerie Longo, Pat Zanzonico, Santosha A. Vardhana

**Author notes:** Current address for Hui Liu: Proteomics Core Laboratory, Memorial Sloan Kettering Cancer Center, New York, NY, USA. Current address for Miguel de Jesus: Vanderbilt Biophotonics Center, Vanderbilt University, Nashville, TN, USA.

## Abstract

Loss of mitochondrial function contributes to CD8^+^ T cell dysfunction during persistent antigen encounter. How chronic antigen leads to this metabolic dysfunction remains unclear. Here, we show that TCR-dependent mitochondrial NADH accumulation drives production of ROS, ultimately leading to mitochondrial dysfunction. Among TCR-dependent proximal signaling components, MEK inhibition uniquely reduced nutrient uptake and mitochondrial NADH accumulation while increasing proliferation. As a result, MEK inhibition during chronic TCR stimulation reduced terminal T cell exhaustion. Mechanistically, we found that chronic MEK activation in T cells drove ATP demand by increasing global protein synthesis rates *in vitro* and *in vivo*. MEK inhibition reversed chronic TCR stimulation-driven increases in RNA polymerase II CTD phosphorylation, reducing transcription rates at effector- and terminal-exhaustion associated genes while maintaining transcription of memory-associated genes. These findings establish MEK-dependent metabolic demand as a driver of T cell exhaustion and elucidate the role of MEK inhibition in enhancing immunotherapy efficacy.

## INTRODUCTION

T cell exhaustion (Texh) is an established barrier to effective immunity against chronic infections and cancer^1^. This dysfunctional state develops in CD8^+^ T cells encountering persistent antigen *in vivo* and is characterized by the upregulation of inhibitory immunoreceptors such as Programmed Death-1 (PD-1), Cytotoxic T-Lymphocyte Antigen 4 (CTLA-4) and Lymphocyte Activation Gene-3 (LAG-3), as well as functional alterations, including reduced proliferative and cytotoxic capacity^1–5^. Over the past twenty years, substantial insights have been made into the molecular drivers of transcriptional and epigenetic alterations present in Texh^6–17^. However, the drivers of functional inactivation of Texh remain incompletely understood.

T cell activation is accompanied by substantial metabolic rewiring to meet the bioenergetic demands of proliferation and cytotoxic function, including a CD28-dependent increase in glucose uptake^18^. Recently, several groups have identified mitochondrial dysfunction as a key contributor to the functional defects of Texh^19–23^, with impaired oxidative phosphorylation and mitochondrial ATP synthesis leading to a loss of proliferative capacity^19^. These mitochondrial defects could be rescued by neutralizing mitochondrial reactive oxygen species (ROS) accumulation, either with the antioxidant N-acetylcysteine, through the overexpression of glutathione peroxidase-4 (Gpx4), or by bolstering NAD^+^ synthesis via nicotinamide riboside supplementation. While sustained TCR signaling is required for Texh generation^4,24–28^, the molecular pathways that initiate and sustain Texh metabolic dysfunction remain poorly defined.

The MEK/ERK signaling cascade is activated downstream of T cell receptor (TCR) engagement^29^ and is required for T cell activation and proliferation^30,31^. Self-limited MEK activation upon initial antigen recognition is essential for optimal T cell responses^32^. However, whether Texh exhibit sustained MEK activation and the consequences of sustained MEK signaling in Texh, remain unexplored. Given its role in regulating the metabolic behavior of other cell types^33^, such as embryonic stem cells^34,35^, the contribution of MEK signaling to the metabolic reprogramming of T cells warrants further investigation. Inhibition of MEK has also been shown to potentiate anti-tumor immunity^36–46^, yet the mechanism by which MEK promotes intratumoral T cell dysfunction remains incompletely understood.

In this study, we establish MEK-dependent protein synthesis as the primary driver of bioenergetic dysfunction in Texh. Sustained TCR signaling was sufficient to increase glucose uptake, mitochondrial NADH generation and mitochondrial ROS accumulation prior to the development of mitochondrial dysfunction, indicating increased ATP demand is an upstream driver of mitochondrial ROS accumulation in T cells. Interrogation of proximal signal transduction components downstream of TCR triggering identified MEK as the central driver of the increased bioenergetic demand that characterizes Texh. Inhibiting MEK signaling in activated T cells was sufficient to attenuate nutrient uptake and mitochondrial ROS accumulation while restoring T cell proliferation. As a result, MEK inhibtion reduced terminal T cell exhaustion both *in vitro* and *in vivo*. Mechanistically, we found that chronic MEK activation is required for global, RNA polymerase II-dependent nascent transcription, and that inhibiting MEK attenuated transcription of genes associated with terminal exhaustion while maintaining transcription of genes associated with self-renewal. These findings highlight MEK-dependent ATP demand as a driver of terminal T cell exhaustion and establish a mechanism by which bioenergetic demand couples TCR signaling to Texh face specification.

## RESULTS

### Chronic TCR engagement drives mitochondrial NADH to support increased ATP demand

Chronic TCR stimulation has been associated with mitochondrial dysfunction and ROS accumulation, contributing to the loss of proliferative capacity that characterizes terminal exhaustion^47^. To determine the proximal impact of chronic TCR stimulation on mitochondrial function, we longitudinally measured mitochondrial activity and ROS accumulation in activated CD8^+^ T cells expanded in the presence or absence of persistent TCR stimulation (Figure 1A). ROS levels in T cells expanded in the presence of cytokine only decreased over time; in contrast, chronic TCR stimulation increased cellular ROS levels within 48 hours and this accumulation continued increasing over time (Figure 1B). ROS is primarily generated within the electron transport chain (ETC) at complexes I and III and has several potential causes, including either an increased rate of electron donation from mitochondrial NADH or an impaired ability to deliver electrons to molecular oxygen at complex IV^48^. Extracellular flux analysis of T cells during either acute or chronic TCR stimulation revealed that oxygen consumption rates (OCR) in chronically stimulated T cells were similar to acutely stimulated T cells after 48 hours but declined after 144 hours of persistent TCR stimulation (Figure 1C).

**Figure 1.**
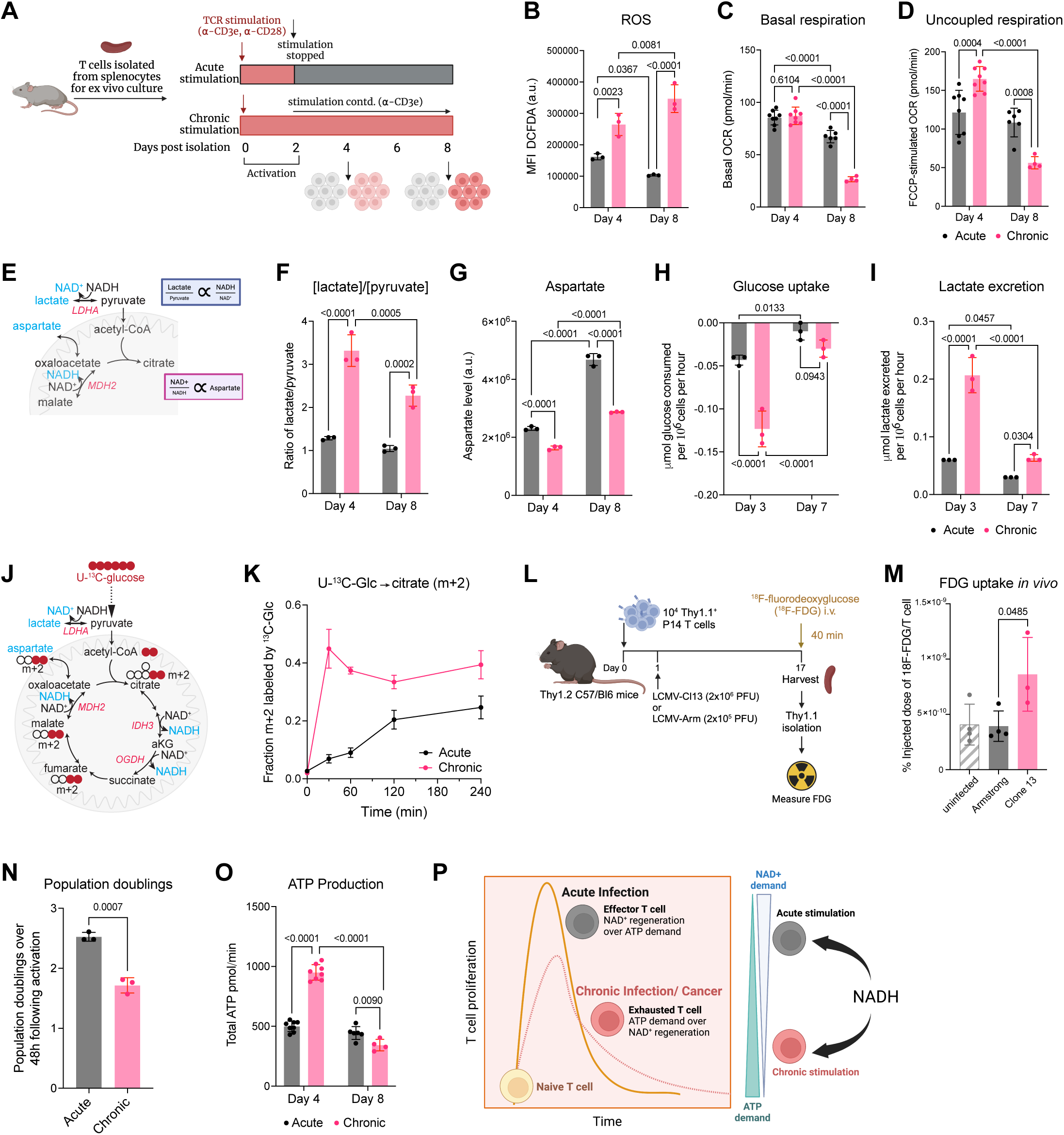
Chronic TCR engagement drives mitochondrial NADH to support increased ATP demand. (A) Experimental strategy for acute and chronic TCR stimulation for temporally resolved metabolomics analyses. (B) ROS levels in acutely (gray) and chronically (pink) stimulated T cells, measured by DCFDA fluorescence (MFI). Data are presented as mean fluorescence intensity (MFI) ± SD (n=3 per group). (C and D) Basal OCR (C) and OCR upon FCCP administration (D) in acutely and chronically stimulated T cells. Error bars are SEM. (E) Schematic of key biochemical pathways highlighting the lactate-to-pyruvate ratio and aspartate pool as proxies for the NADH/NAD⁺ redox state. (F and G) Quantification of the lactate-to-pyruvate ratio (F) and aspartate pool size (G). (H and I) Quantification of median glucose consumption (H), and lactate production (I) at 24 and 120 hours of chronic stimulation, compared to acute stimulation. (J) Schematic illustrating ¹³C-labeling patterns in TCA cycle metabolites following U-¹³C-glucose metabolism after the first round. Colored circles represent 13C, empty circles represent 12C. (K) Time course of m+2 citrate labeling in T cells treated with U-¹³C-glucose for 30, 60, 120, and 240 minutes. Two-way ANOVA with Sidak’s multiple comparisons test was used (p< 0.001 at t > 0). (L and M) Experimental strategy (L) and quantification of ¹⁸F-FDG uptake (M) in antigen-specific P14 CD8⁺ T cells from the spleen of mice infected with LCMV Armstrong or LCMV Clone 13, or uninfected controls. Data are presented as the mean ± SD of n =12 per group. The experiment was independently repeated twice. (N) Number of population doublings after 48 hours of culture in acutely or chronically stimulated T cells. Statistical significance was determined using one-way ANOVA. (O) Total ATP production calculated from Extracellular Flux Analysis including contributions from both glycolysis and mitochondrial respiration. Error bars are SEM. (P) Schematic illustration highlighting relative prioritization of NAD^+^ regeneration versus ATP synthesis in T cells following acute (gray) versus chronic stimulation (red). Data are presented as mean ± SD (n = 3–8 per group) from representative experiments independently repeated at least three times. Statistical significance was determined using two-way ANOVA unless stated otherwise (p-values indicated).

This indicates that the observed ROS accumulation at 48 hours occurred in the absence of an observable loss in ETC function, suggesting that increased mitochondrial NADH drives ROS accumulation in chronically stimulated T cells. Further supporting this, ETC-uncoupled OCR (as measured by OCR in cells treated with the uncoupler carbonyl cyanide-4-(trifluoromethoxy)phenylhydrazone (FCCP)) was significantly increased in chronically stimulated T cells, indicative of increased mitochondrial NADH generation in excess of the rate of ADP-coupled respiration (Figure 1D, S1A). Mitochondrial NADH levels are reflected by both the ratio of intracellular lactate to pyruvate^49–51^ as well as steady state levels of intracellular aspartate^52,53^ (Figure 1E). We observed an increase in lactate:pyruvate ratio (Figure 1F) and a decrease in intracellular aspartate pools (Figure 1G) within 48 hours of chronic TCR stimulation, consistent with increased mitochondrial NADH levels.

To identify the mechanism by which TCR stimulation supports mitochondrial NADH generation, we measured glucose uptake and metabolism in T cells in response to chronic TCR stimulation. 48 hours of chronic TCR stimulation was sufficient to increase glucose uptake (Figure 1H) and lactate production (Figure 1I, S1B) as well as entry of glucose-derived carbons into the TCA cycle (Figure 1J-K). These results further confirm that the increased mitochondrial NADH levels in chronically stimulated T cells is driven by NADH generation rather than by impaired NADH clearance. To confirm that chronic TCR stimulation increases glucose uptake *in vivo*, we measured uptake of ^18^F-fluorodeoxyglucose (FDG) in activated T cells undergoing chronic TCR stimulation in mice infected with LCMV-Clone 13^54^ (Figure 1L). FDG levels were higher in T cells isolated from LCMV-Clone 13 infected mice compared to T cells from resolving LCMV-Armstrong infection (Figure 1M). Consistent with mitochondrial NADH accumulation, as prior reports have shown that NAD^+^ regeneration is limiting for cellular proliferation, including in primary T cells^55^, we found that chronic TCR stimulation immediately reduced T cell proliferation (Figure 1N). Despite this, the calculated ATP production by T cells under chronic TCR stimulation was increased (Figure 1O, S1C), suggesting that chronic TCR stimulation-dependent glucose uptake and mitochondrial NADH generation supports ATP production at the expense of proliferative capacity (Figure 1P). Taken together, these results indicate that persistent TCR stimulation increases nutrient uptake and mitochondrial NADH generation to support increased ATP demand.

### TCR-dependent bioenergetic dysfunction requires persistent MEK activation

We next asked whether our observed increase in bioenergetic demand and mitochondrial ROS during chronic TCR stimulation depended on the activity of a specific TCR-dependent proximal signaling pathway. We therefore performed a small molecular inhibitor screen targeting proximal TCR-dependent signal transduction molecules in activated T cells cultured with persistent TCR stimulation (Figure 2A). Of the inhibitors tested, MEK inhibition (MEKi) uniquely reduced both the calculated rate of ATP synthesis (Figure 2B, S2A-S2C) as well as ROS accumulation (Figure 2C) during persistent TCR stimulation. We confirmed that chronic antigen exposure was sufficient to increase MEK/Erk activation, both *in vitro* after 48 hours of persistent stimulation (Figure 2D) and *in vivo* in T cells from LCMV-Clone 13 infection (Figure 2E-F). Moreover, direct measurements of glucose uptake (Figure 2G) and lactate excretion (Figure 2H) revealed that MEKi reduced glucose consumption and glycolytic activity in chronically stimulated T cells, indicating that inhibiting MEK diminishes the bioenergetic demand induced by chronic stimulation. Although MEK was not the only proximal TCR signaling component whose inhibition reduced glucose uptake, only MEKi reduced glucose uptake while increasing proliferation in chronically stimulated T cells (Figure 2I-J, S2D-S2J). This suggests that MEKi reverses mitochondrial NADH accumulation during chronic TCR stimulation. Consistent with this hypothesis, MEKi during chronic TCR stimulation decreased the lactate:pyruvate ratio (Figure 2K) and increased intracellular aspartate pools (Figure 2L). These results indicate that MEK activation during chronic TCR stimulation drives nutrient uptake, mitochondrial NADH accumulation, and loss of T cell proliferation.

**Figure 2.**
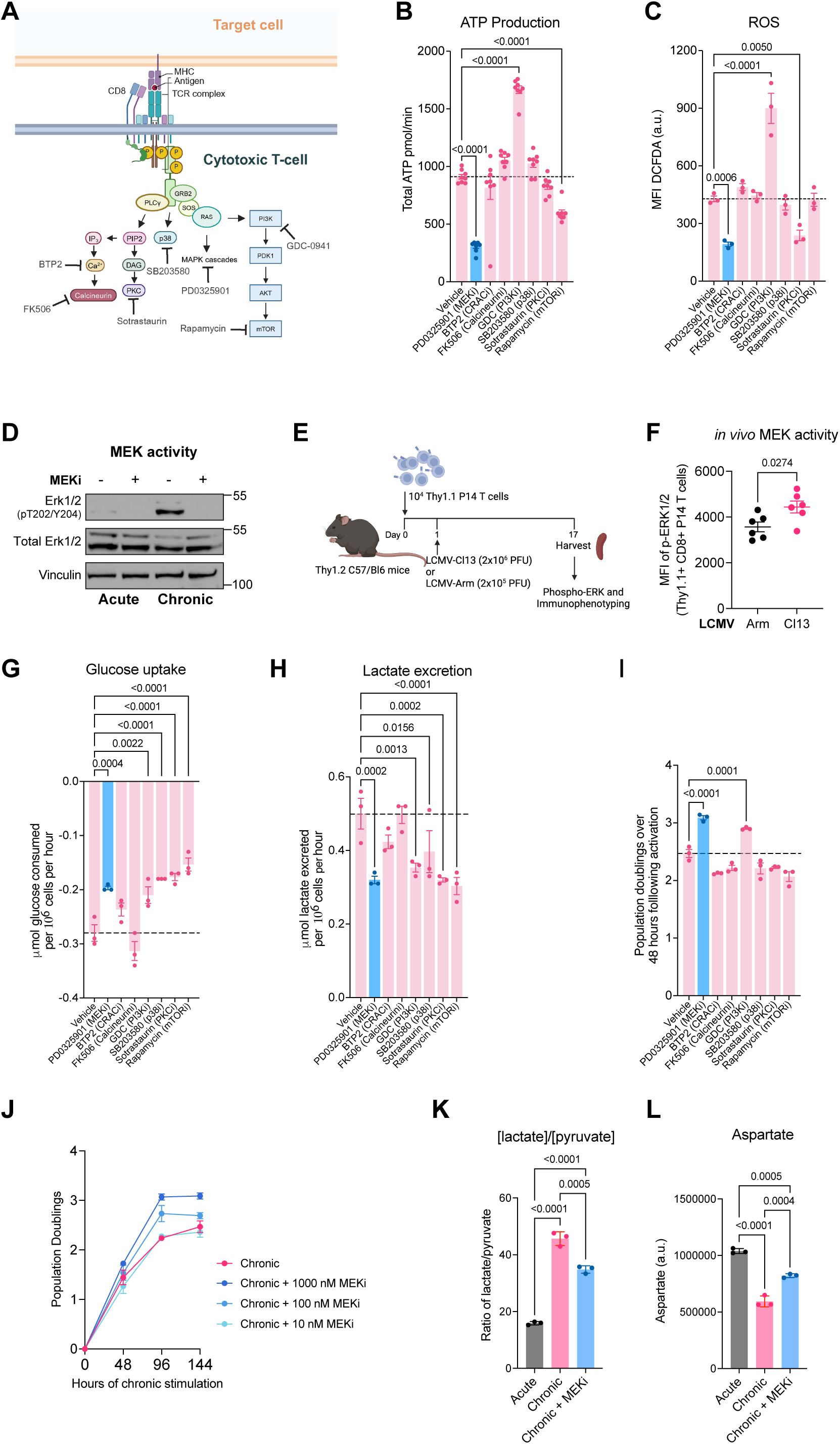
TCR-dependent bioenergetic dysfunction requires persistent MEK activation. (A) Schematic of TCR proximal signaling pathways, highlighting key intermediates, and corresponding inhibitors. (B and C) Total ATP production (calculated from glycolysis and mitochondrial respiration) (B), and ROS levels (measured by DCFDA fluorescence, MFI) (C) in T cells treated with indicated inhibitors or vehicle during 48 hours of chronic TCR stimulation. (D) Immunoblot showing total and T202/Y204 phosphorylated Erk1/2 in acutely and chronically stimulated T cells, with Vinculin as a loading control. Western blot was performed two independent times. Uncropped blots are available as source data. (E and F) Experimental strategy (E) and quantification of Erk1/2 phosphorylation (F) in antigen-specific P14 CD8⁺ T cells from the spleen of LCMV-infected mice. The experiment was independently repeated twice. (G and H) Quantification of median glucose uptake (G) and lactate excretion (H) in T cells treated with indicated inhibitors or vehicle during 48 hours of chronic stimulation. (I) Number of population doublings in chronically stimulated T cells treated with indicated inhibitors or vehicle for 48 hours. (J) Population doublings of chronically stimulated T cells with or without MEK inhibitor treatment at specified concentrations over 6 days of chronic TCR stimulation, p<0.0001 for 1000 nM, p= 0.0218 for 100 nM, p= 0.5378 for 10 nM. (K and L) Quantification of the lactate-to-pyruvate ratio (K) and aspartate pool size (L) in T cells stimulated under the indicated conditions for 48 hours. Data are presented as mean ± SD (n = 3–8 per group) from representative experiments independently repeated at least three times. Statistical significance was determined using one-way ANOVA (p-values indicated). The dotted line represents the mean value for T cells treated with vehicle.

### MEKi decreases terminal T cell exhaustion and enhances persistence in vivo

Terminal T cell exhaustion is marked by a loss of proliferative capacity. Our finding that chronic TCR-driven MEK activation leads to mitochondrial NADH accumulation and impaired proliferation suggested that MEKi might be sufficient to limit terminal T cell exhaustion induced by chronic antigen stimulation. Indeed, MEKi restored the proliferation of chronically stimulated T cells while having no notable effect on the proliferation of acutely stimulated T cells (Figure 3A). To determine the impact of MEKi on chronic antigen-driven activation of the T cell exhaustion program, we analyzed T cells that were acutely or chronically stimulated in the presence or absence of MEKi using both flow cytometry and bulk RNA sequencing. MEKi did not reduce chronic antigen-driven upregulation of PD-1, suggesting that chronic MEK activation is not required for all aspects of the T cell exhaustion program (Figure 3B). However, MEKi reduced expression of inhibitory receptors enriched in terminally exhausted T cells, such as Tim-3 and LAG-3, while restoring expression of progenitor-associated markers, such as SlamF6^56^ (Figure 3B). These results suggest that chronic MEK activation drives the transition of exhausted T cells from a progenitor-like to a terminally exhausted state. Bulk RNA sequencing analysis of chronically stimulated MEKi-treated T cells revealed upregulation of genes enriched in Texh with a progenitor phenotype and downregulation of genes enriched in Texh with a terminally exhausted phenotype^47^ (Figure 3C). This MEKi-driven attenuation of terminal Texh differentiation may be due in part to a reduction in Calcium-dependent NFAT activity, which is a key transcriptional driver of the Texh state^8^. Consistent with this hypothesis, MEKi suppressed expression of NFAT target genes (Figure S3A). Nuclear NFAT activity requires both intracellular calcium and mitochondrial ROS; MEKi did not alter intracellular calcium dynamics in chronically stimulated T cells (Figure 3D), indicating that MEK likely promotes NFAT activity by increasing mitochondrial ROS. The role of MEK activation in driving terminal T cell exhaustion was both dose-dependent and continuous, as we observed both, a dose- and time-dependent effect, of MEKi on T cell proliferation and expression of terminal exhaustion markers during chronic TCR stimulation (Figure S3B-C). This suggests that the extent of MEK activation during chronic TCR stimulation acts as a metabolic rheostat that determines the balance between progenitor-like and terminally differentiated states in response to antigen dose and persistence.

**Figure 3.**
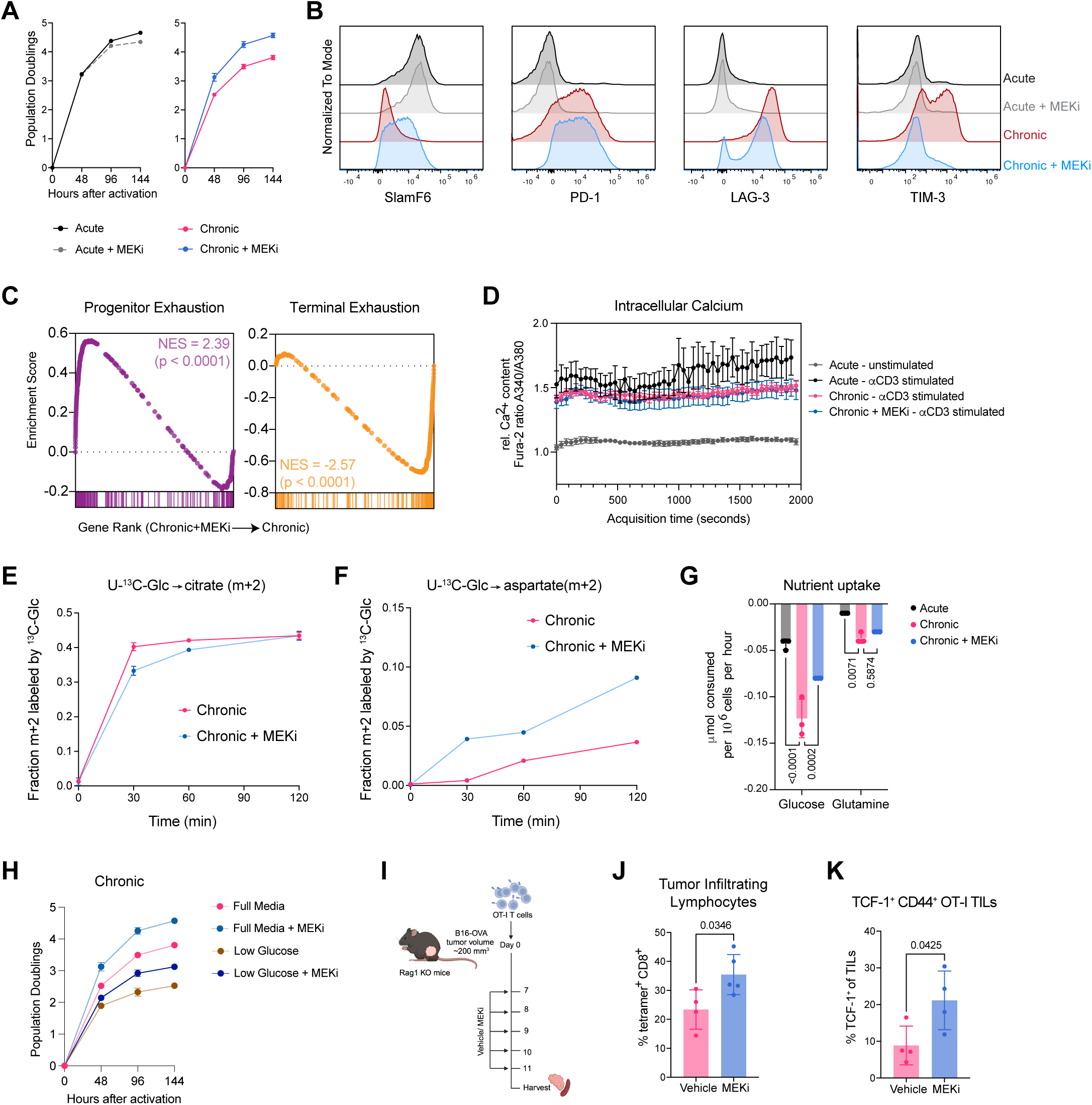
MEKi decreases terminal T cell exhaustion and enhances persistence *in vivo*. (A) Population doublings of acutely or chronically stimulated T cells with or without MEKi (100 nM PD0325901) treatment over time, p<0.0001 from 48 hours for chronic. (B) Expression of exhaustion markers (SlamF6, PD-1, LAG-3, TIM-3) in acute and chronic T cells with or without MEKi treatment. (C) Gene set enrichment analysis (GSEA) of genes enriched in terminal and progenitor subsets of exhausted tumor-infiltrating T cells^47^ in T cells cultured with or without MEKi treatment during 6 days of chronic TCR stimulation. Normalized enrichment scores (NES) and p-values are indicated. (D) Calcium flux measurements of T cells under chronic stimulation with or without MEKi treatment, assessed by Fura-2 ratiometric imaging. T cells activated under acute conditions exhibit calcium flux upon αCD3 stimulation (black), whereas no calcium flux is detected in these cells without αCD3 stimulation (gray). Error bars are SEM. The experiment was independently repeated twice. (E and F) Time course of m+2 labeling of citrate (E) and aspartate (F) by 13C-glucose in chronically stimulated T cultured cells with or without MEKi. Two-way ANOVA with Sidak’s multiple comparisons test was used (p< 0.001 at t= 30 min). (G) Glucose and glutamine uptake by T cells cultured in the presence of acute or chronic stimulation with or without MEKi treatment. (H) Population doublings of T cells cultured during chronic stimulation in the presence or absence of a MEK inhibitor in RPMI containing full (10 mM) or reduced (Low, 1 mM) glucose availability. (p<0.0001) (I) Experimental design to assess the impact of MEK inhibition on intratumoral T cell abundance *in vivo*. (J and K) Quantification of tetramer⁺ CD8⁺ tumor-infiltrating lymphocytes (TILs) (J) and TCF-1⁺ CD44⁺ OT-I TILs (K) in Trametinib- and vehicle-treated groups. Statistical significance was determined by unpaired t-test (p-values indicated). Statistical significance was determined by unpaired t-test (p-values indicated). Data are presented as mean ± SD (n = 3–8 per group) from representative experiments independently repeated at least three times. Statistical significance was determined using two-way ANOVA unless stated otherwise (p-values indicated).

To understand how MEK inhibition during chronic TCR stimulation enables sustained T cell proliferation while simultaneously reducing metabolic demand, we performed isotopologue tracing using uniformly labeled 13C-glucose (U-13C-glc)^57^. Analysis of TCA cycle flux revealed that MEKi reduced the rate of m+2 citrate labeling from glucose, consistent with decreased flux of glucose-derived carbons into the TCA cycle (Figure 3E). However, m+2 malate labeling remained comparable with or without MEKi (Figure S3D). Malate can be generated via NAD^+^-dependent oxidative decarboxylation of citrate within the TCA cycle or via ATP citrate lyase-mediated generation of cytosolic oxaloacetate via a recently described non-canonical TCA cycle^58^ (Figure S3E). The relative engagement of canonical to non-canonical TCA cycle activity can be measured by the ratio of m+2 malate relative to m+2 citrate. This ratio was increased with MEKi (Figure S3F). Similarly, m+2 labeling of aspartate increased with MEKi (Figure 3F), consistent with an increased rate of oxidative decarboxylation of glucose-derived citrate within the TCA cycle. As multiple enzymatic reactions within the TCA cycle require NAD^+^ as a co-factor, these results are consistent with MEKi enhancing T cell proliferation during chronic TCR stimulation by restoring mitochondrial NAD^+^, thereby enabling oxidative nucleotide metabolism.

Given that MEKi relieved TCR-driven bioenergetic demand and nutrient uptake, we hypothesized that MEKi might enable T cell persistence in nutrient-depleted environments, including within tumors. We confirmed that MEKi was sufficient to reduce both glucose and glutamine uptake during chronic TCR stimulation (Figure 3G). We subsequently cultured T cells under 10-fold reduced availability of glucose (1 mM) or glutamine (0.2 mM) during acute or chronic TCR stimulation. While acutely stimulated T cells exhibited only a ∼6% decrease in proliferation (Figure S3G), chronically stimulated T cells were more sensitive to glucose depletion, showing a ∼33% reduction in proliferation (Figure 3H). MEKi restored the proliferative capacity of these T cells in both glucose-replete and glucose-limited conditions (Figure 3H). Similar results were observed when T cells were cultured in media with reduced extracellular glutamine concentrations (Figure S3G-H), indicating that MEKi broadly improves T cell proliferation during nutrient restriction. Based on these findings, we hypothesized that MEKi-treated, antigen-specific T cells would have enhanced intratumoral persistence. To test this, we adoptively transferred naive CD8^+^ TCR transgenic OT-I T cells into B16-OVA-bearing mice followed by treatment with either vehicle or MEKi (Trametinib, 1mg/kg) for 5 days (Figure 3I). MEKi-treated mice contained approximately double the intratumoral abundance of antigen-experienced OT-I cells compared to controls (Figure 3J-K).

### Bioenergetic demand during chronic TCR stimulation is driven largely by MEK-driven protein synthesis

We next sought to identify the underlying driver of TCR driven, MEK-dependent ATP demand in chronically stimulated T cells. Gene set enrichment analysis of transcriptomes from chronically stimulated T cells cultured in the presence or absence of MEKi following T cell activation revealed that the most downregulated gene sets were associated with protein synthesis, suggesting that TCR-dependent MEK activation increases mRNA translation rates during chronic antigen encounter (Figure 4A). Since protein synthesis accounts for a significant fraction of ATP consumption in metabolically active cells^59^ due to the requirement of GTP for both tRNA charging and amino acid transfer to elongating peptide strands, we hypothesized that the increased ATP demand observed in chronically stimulated T cells is driven by increased rates of protein synthesis. Consistent with this hypothesis, both cell volume, which generally reflects cell mass and intracellular protein content, as well as mTORC1 activity, which is a known regulator of cell growth and protein synthesis, were increased in activated T cells undergoing chronic TCR stimulation in a MEK-dependent manner (Figure S4A and Figure 4B, respectively). Increased protein synthesis can also trigger ER stress^60^, and this has been reported in exhausted T cells *in vivo*^61^. We found that 48 hours of persistent TCR stimulation was sufficient to activate components of the ER stress response, including stabilization of ATF4 and Binding Immunoglobulin Protein (BiP), both of which were reversed by MEKi (Figure S4B).

**Figure 4.**
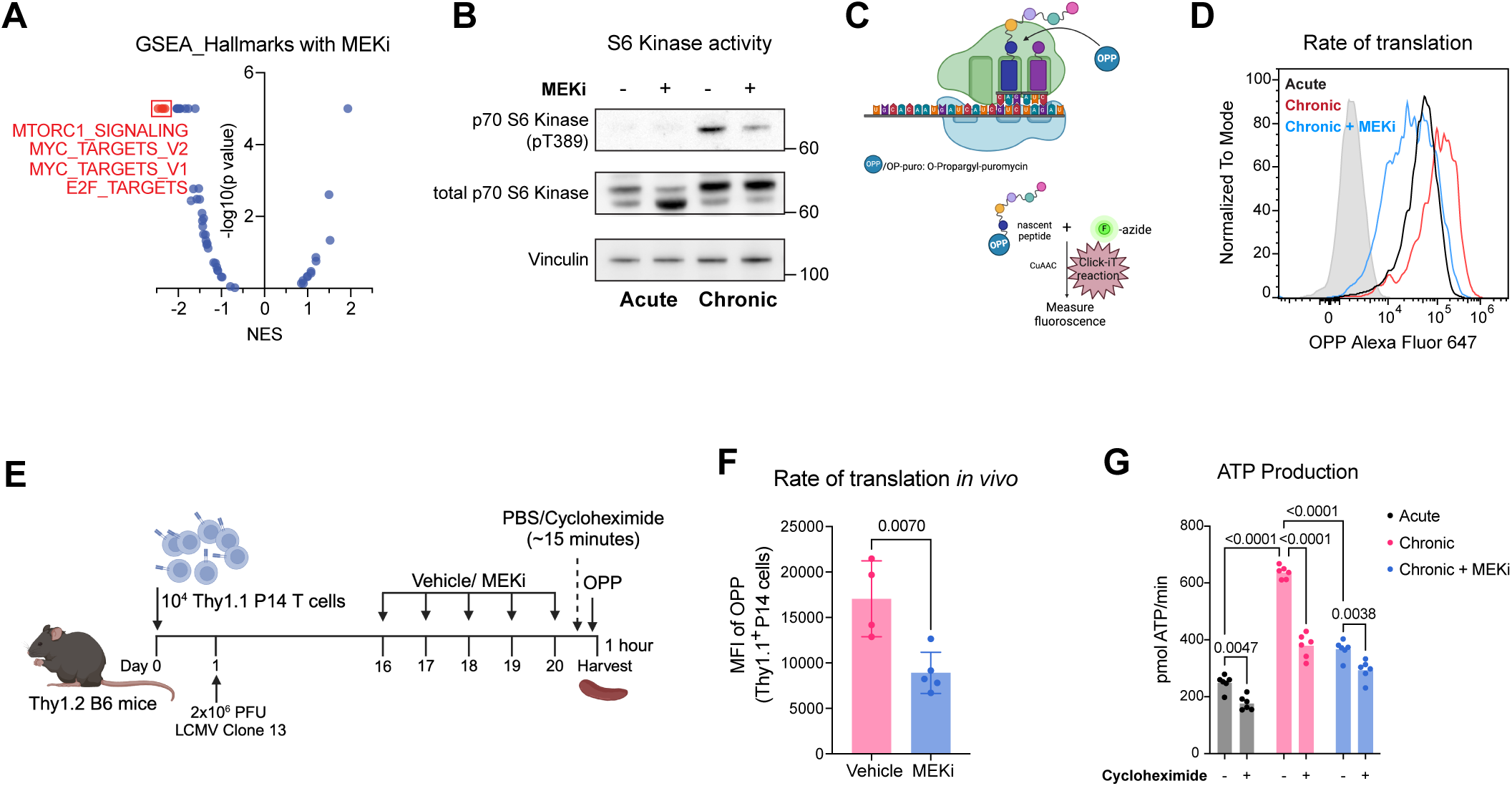
Bioenergetic demand during chronic TCR stimulation is driven largely by MEK-driven protein synthesis. (A) Gene Set Enrichment Analysis (GSEA) of hallmark pathways affected by MEKi in chronically stimulated T cells, showing NES (Chronic+MEKi vs. Chronic) on the x-axis and statistical significance (-log10 p-value) on the y-axis. (B) Immunoblot analysis of total and pT389 phosphorylated p70 S6 kinase in acute and chronically stimulated T cells, with or without MEKi treatment. Vinculin serves as a loading control. (C) Schematic representation of an O-propargyl puromycin (OPP) assay used to measure single cell rates of nascent protein synthesis. OPP incorporation into nascent peptides allows fluorescence-based quantification via a Click-IT reaction. (D) Representative histogram of OPP incorporation by acutely or chronically stimulated T cells with or without MEKi treatment. (E) Experimental setup to measure the impact of MEK inhibition on T cell-specific translation rates during chronic LCMV infection *in vivo*. (F) Quantification of OPP incorporation in Thy1.1⁺ P14 T cells, measured as Mean Fluorescence Intensity (MFI). Statistical significance was determined by unpaired t-test (p-value indicated). (G) Nascent protein synthesis-dependent ATP production rates in acute, chronic, and MEKi-treated chronic T cells following acute treatment with either cycloheximide or PBS control. Statistical significance was determined using two-way ANOVA (p-values indicated).

To directly measure protein synthesis rates, we labeled nascent proteins with O-propargyl-puromycin (OPP), which is incorporated non-specifically into elongating peptide strands and can be conjugated to an azide-linked fluorophore via click chemistry (Figure 4C)^62^. We found that chronic TCR stimulation significantly increased OPP fluorescence, and this effect was attenuated by MEKi (Figure 4D). To confirm that TCR-dependent MEK activation regulates T cell protein synthesis rates *in vivo*, we adoptively transferred naive CD8^+^ TCR transgenic P14 T cells into mice infected with chronic LCMV-Clone 13 infection, followed by treatment with either vehicle or a MEK inhibitor (Trametinib, 1mg/kg) for 5 days. One hour prior to being sacrificed, mice were administered OPP (49.5 mg/kg i.p.)^63^ with or without cycloheximide (CHX) pre-treatment, followed by rapid harvest, cell surface labeling, and click chemistry-based fluorophore conjugation (Figure 4E). P14 T cells from MEKi-treated LCMV-Clone 13 infected mice showed reduced expression of terminal T cell exhaustion markers (Tim-3) and increased expression of memory-associated markers (SlamF6) (Figure S4C) as well as reduced rates of OPP incorporation, consistent with a MEK-dependent increase in protein synthesis rates during chronic TCR stimulation *in vivo* (Figure 4F). Finally, we asked whether a chronic TCR stimulation driven increase in protein synthesis was sufficient to increase ATP demand in T cells. To test this hypothesis, we measured the impact of CHX treatment on calculated ATP consumption rates based on extracellular flux analysis in chronically stimulated T cells. Inhibition of translation with CHX rapidly reduced ATP consumption rates in chronically stimulated T cells, confirming that protein synthesis is a major net consumer of ATP during chronic TCR stimulation (Figure 4G, S4D).

### TCR-dependent MEK activation increases nascent transcription of genes associated with terminal exhaustion

CD8^+^ T cell exhaustion is characterized by the sequential activation of transcriptional programs that are associated with progenitor-like and terminally differentiated states^47,64–67^. In eukaryotic cells, mRNA synthesis is primarily driven by RNA Polymerase II (RNAPII), which is recruited to gene promoters by general transcription factors, such as TFIIH, as part of the transcription initiation complex. The activity of RNAPII is regulated at the levels of both transcription initiation and elongation through phosphorylation of serines S5 and S2 within the C-terminal domain (CTD), respectively^68^. Since in embryonic stem cells, CTD phosphorylation of RNAPII is sensitive to MEKi^35^, we hypothesized that our observed TCR-driven, MEK-dependent increase in protein synthesis rates might be secondary to increased Pol II activity. We found that chronic TCR stimulation was sufficient to increase RNAPII CTD phosphorylation at both S5 and S2 in a manner that was reversed by MEKi (Figure 5A), suggesting that TCR-dependent MEK activation increases global transcription initiation and elongation. To test this directly, we assessed nascent transcription during chronic TCR stimulation by pulsing T cells with 5-ethinyluridine (5EU), a cell-permeable uridine analog that incorporates into newly synthesized mRNA and can be analyzed via click chemistry-based conjugation to either a fluorophore for nascent mRNA quantitation, or biotin for streptavidin-based mRNA isolation, cDNA library generation, and Illumina sequencing of nascent mRNA transcripts (Figure 5B)^69^. Consistent with increased RNAPII CTD phosphorylation, we found that chronic TCR stimulation led to increased nascent RNA synthesis rates in a MEK-dependent manner (Figure 5C).

**Figure 5.**
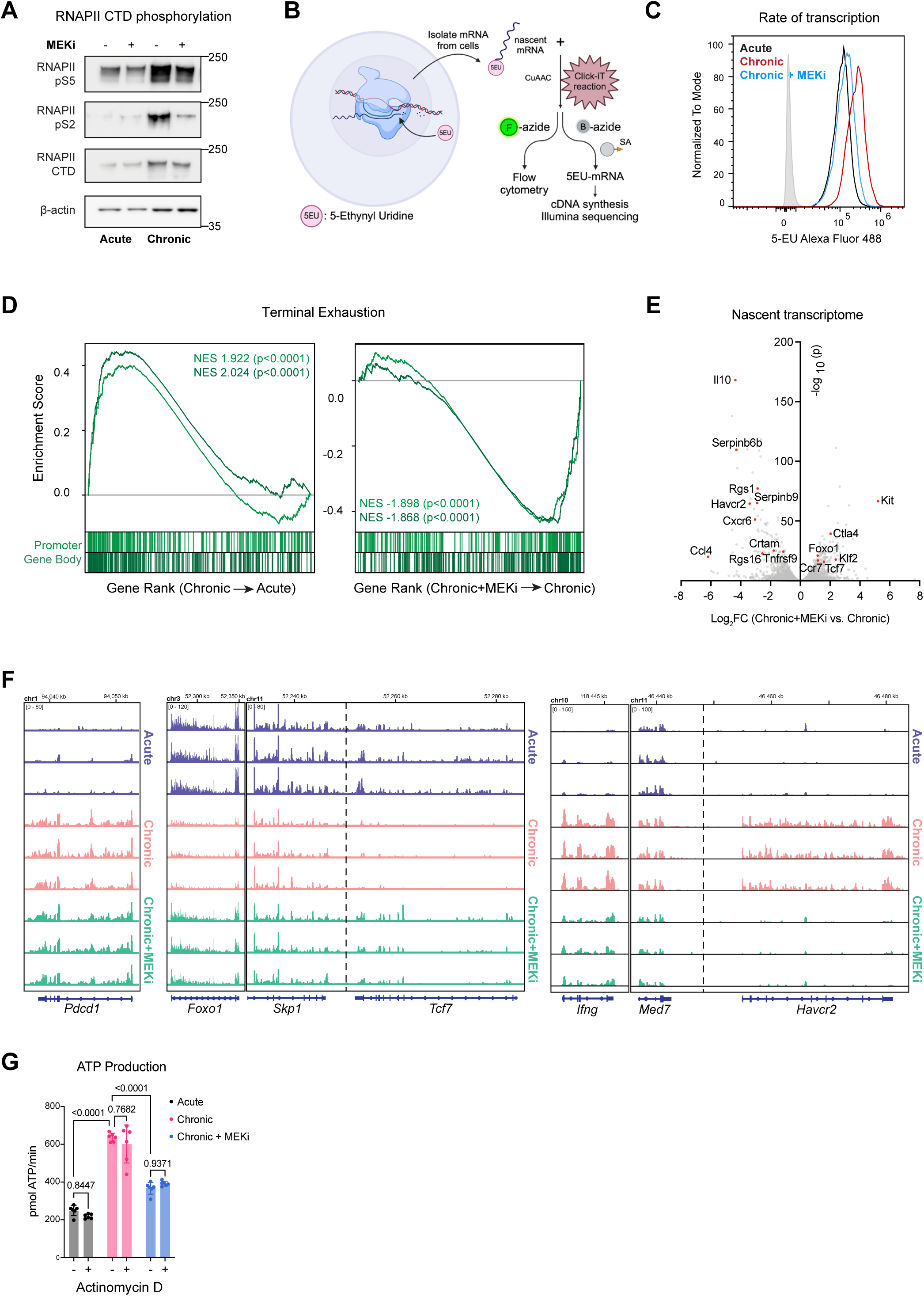
TCR-dependent MEK activation increases nascent transcription of genes associated with terminal T cell exhaustion. (A) Immunoblot analysis of phosphorylated (pS5 and pS2) and total RNA Polymerase II CTD in acutely and chronically stimulated T cells with or without MEKi treatment. β-actin serves as a loading control. (B) Schematic representation of 5-Ethynyluridine (5-EU) assay used to measure rate of transcription. 5-EU is incorporated into nascent RNA, and the labeled RNA can be detected by flow cytometry using a fluorophore-conjugated azide. Additionally, 5-EU-labeled mRNA can be isolated via a Biotin-azide Click-IT reaction, followed by streptavidin-based capture and sequencing for further analysis. (C) Representative histogram of 5-EU incorporation in acute, chronic, and MEKi-treated chronic T cells. (D) Gene Set Enrichment Analysis (GSEA) of genes enriched in terminally exhausted T cells^47^ in chronic T cells compared to acute T cells (left) and in chronic T cells with or without MEKi treatment (right). Normalized enrichment scores (NES) and p-values are indicated. (E) Volcano plot of differentially expressed nascent transcripts in chronic T cells treated with MEKi, showing log_2_ fold-change (Chronic+MEKi vs. Chronic) on the x-axis and statistical significance (-log_10_ p-value) on the y-axis. Select genes are highlighted. (F) Genome browser tracks depicting nascent RNA transcription levels in acute, chronic, and chronic + MEKi T cells. Tracks represent 5-EU labeled nascent RNA. (G) Nascent transcription-dependent ATP production rates in acute, chronic, and MEKi-treated chronic T cells following acute treatment with either actinomycin D or PBS control. Statistical significance was determined using two-way ANOVA (p-values indicated).

Finally, to determine the gene-specific impact of TCR-dependent MEK activation on transcription, we analyzed nascently synthesized mRNA transcripts from T cells undergoing acute or chronic TCR stimulation *in vitro,* in the presence or absence of MEKi. We found that chronic TCR stimulation increased nascent synthesis of mRNAs known to be upregulated in terminally exhausted T cells, including genes encoding inhibitory receptors such as *Havcr2* and *Lag3,* as well as genes encoding cytotoxic proteins such as *Ifng* and *Gzmb* (Figure 5D-F). In contrast, MEK inhibitor-treated T cells reduced nascent transcription of genes associated with terminal T-cell exhaustion while increasing nascent transcription of memory associated genes such as *Sell*, *Foxo1* and *Tcf7* (Figure 5E-F). These results are consistent with MEK-dependent transcription and translation of terminal exhaustion-associated genes as a bioenergetic driver of T cell exhaustion during chronic TCR stimulation. However, pre-treatment of chronically stimulated T cells with Actinomycin D for 12 hours, which is sufficient to suppress nascent transcription, did not alter ATP consumption rates, suggesting that mRNA translation is the primary source of increased ATP consumption in response to TCR-dependent MEK activation (Figure 5G).

## DISCUSSION

In this study, we identify MEK activation as a primary driver of chronic antigen-driven T cell metabolic dysfunction and terminal differentiation. The molecular drivers of terminal CD8^+^ T cell differentiation have been of increasing interest in recent years, given their predictive role in immunotherapy outcomes^47,67,70^. Most distinct cell types in multicellular organisms are maintained by a pool of self-renewing progenitor cells that give rise to differentiated sub-populations with tissue-specific functions. Exhausted T cells fit within this framework, with a pool of TCF-1^+^ progenitor-like cells that prioritize self-renewal, and a population of terminally exhausted cells dedicated to target-cell elimination^47^. Our findings establish that MEK controls the utilization of nutrients to direct these distinct cell fates. During chronic TCR signaling, persistent MEK activation increases RNA Polymerase II-dependent gene transcription and mRNA translation alongside increased nutrient uptake to supply ATP to meet this increased bioenergetic demand. This metabolic switch enables effector differentiation and transient cytotoxic function. However, persistent MEK-dependent mitochondrial NADH generation ultimately drives accumulation of ROS, leading to mitochondrial dysfunction, uncoupling between increased bioenergetic demand and mitochondrial ATP synthesis, and terminal T cell exhaustion. These findings refine current models of T cell exhaustion by positioning MEK as a central regulator of mitochondrial dysfunction and provide a mechanistic rationale for the observed clinical benefits of MEK inhibition in enhancing T cell anti-tumor immunity.

Specifically, MEK inhibition reprograms exhausted T cells by lowering ATP demand through suppression of protein synthesis, thereby promoting T cell persistence in nutrient-deprived tumor microenvironment. Recent studies suggest that pulsatile MEKi, in combination with checkpoint blockade, enhances anti-tumor immunity in mouse models^42^. This aligns with our findings: MEK inhibition reduces metabolic demand and enhances T cell persistence, while checkpoint blockade increases nutrient uptake and cytotoxicity^71^. A rationally designed cyclical regimen of MEK inhibitors and checkpoint blockade could therefore balance T cell persistence with effector function, optimizing therapeutic efficacy. Dynamic monitoring of metabolic parameters in T cells during pulsatile versus continuous MEKi may help refine optimal dosing regimens and improve clinical outcomes.

The allocation of NADH either toward ATP production via oxidative phosphorylation (OXPHOS) or toward NAD^+^ regeneration through glycolysis enables effector versus memory lineage commitments in naive as well as exhausted T cells. MEK activation is a logical candidate to mediate this fate decision as it marks the transition between analog and digital signal output in response to a graded input in lymphoid cells^72^. This is due to expression of a GTP exchange factor, SOS1, whose catalytic activity is amplified by allosteric binding of Ras-GTP^73,74^. This positive feedback loop enables conversion of a continuous signal input to a decisive, digital output, which is essential for naive T cells. An interesting property of this feedback loop is that the decay in loss of allosteric activation of SOS1 enables cells to respond maximally when rechallenged with a comparatively weaker stimulus. This property is essential to allow naive T cells to integrate antigenic signals while migrating between dendritic cells within the draining lymph node, as has been previously described^75–77^. However, under conditions of sustained antigen burden, this feedback loop causes cells to exhibit maximal MEK activation even when antigenic stimulation is discontinuous. Given the role of TCR signal strength in exhaustion, future studies should explore whether modulating MEK activity, either by targeting Sos1 or other upstream regulators of MEK activation, can enhance the durability of T cells expressing high-avidity antigen receptors, including CAR-T cells.

Finally, our observation that MEK functions as a rheostat to titrate ATP production and NAD^+^ regeneration in response to growth factor-dependent demand in T cells raises the question of whether MEK plays a similar role more broadly within cell biology. In embryonic stem cells, MEK inhibition maintains cells in a state of pluripotency, and this is similarly associated with reduced rates of transcription, particularly of lineage-specific genes^35^. Similarly, persistent MEK activation is implicated in senescence of both transformed and non-transformed cells^78^. Future studies are needed to determine whether MEK-dependent modulation of NAD^+^ generation via reduction of pyruvate to lactate and mitochondrial ATP production can be a therapeutic target across cell states.

In summary, our findings establish MEK as a pivotal metabolic checkpoint in T cell exhaustion. By linking chronic antigenic stimulation to mitochondrial dysfunction, our study not only deepens our mechanistic understanding of exhaustion but also lays the groundwork for innovative therapeutic strategies aimed at reprogramming T cell metabolism to enhance immunotherapy.

## RESOURCE AVAILABILITY

### Lead contact

Correspondence and requests for materials should be addressed to the lead contact, Dr Santosha Vardhana.

### Materials availability

This study did not generate new unique reagents.

### Data and code availability

- Nascent RNA-seq data have been deposited at GEO at GEO **<u>GSE293649</u>** and are publicly available as of the date of publication.
- Any additional information required to reanalyze the data reported in this paper is available from the lead contact upon request.

## ACKNOWLEDGMENTS

S.A.V was supported by an NCI K08 Career Development Award (NCI K08 CA237731), a Burroughs Wellcome Fund Career Award for Medical Scientists, a V Foundation Scholar Award, and the Josie Robertson Investigators Program. This work was additional supported by a Cancer Center Support Grant (P30 CA008748). T.M. was supported by the Dorris J. Hutchison Pre-Doctoral Fellowship by Memorial Sloan Kettering. We acknowledge the Integrated Genomics Operation Core at Memorial Sloan Kettering for helping with the RNA sequencing experiments, and the Nuclear Pharmacy of the Molecular Imaging and Therapy Service (Department of Radiology) at Memorial Sloan Kettering for the generous provision of ^18^F-FDG (Sofie, Totowa, NJ) doses for our animal experiments. We are grateful to Michael Bale and Steven Josefowicz for helpful insights on nascent transcript analysis. Finally, we appreciate the members of the Vardhana, Thompson, Finley, and Intlekofer laboratories for their valuable input and discussions.

## AUTHOR CONTRIBUTIONS

T.M. and S.A.V. designed the study and wrote the manuscript. T.M. performed all the experiments. J.R. performed all the computational analysis. M.H. helped with generating independent replicates of the screening experiments. R.R., H.L., T.H., and J.C. contributed to metabolomics experiments. M.D.J. and M.H designed and assisted with Calcium flux experiments. P.Z. and V.L. helped with radioactive experiments. All co-authors reviewed the manuscript prior to submission.

## SUPPLEMENTAL INFORMATION

**Figure S1 Increased metabolic demand within 48 hours of chronic TCR stimulation**

(A) Oxygen consumption rate of T cells following 48 hours (left) or 144 hours (right) of chronic TCR stimulation at baseline or in the presence of ATP synthase inhibition (Oligo), uncoupling agents (FCCP) or complex III/IV inhibition (Rot/AA).

(B) Median lactate excreted per molecule of glucose consumed in acutely and chronically stimulated T cells following initial stimulation, p-value indicated.

(C) Extracellular acidification rate of T cells following 48 hours (left) or 144 hours (right) of chronic TCR stimulation at baseline or in the presence of ATP synthase inhibition (Oligo), uncoupling agents (FCCP) or complex III/IV inhibition (Rot/AA).

**Figure S2 MEK inhibition attenuates T cell metabolic demand while increasing proliferation**

(A and B) Basal OCR (A) and basal ECAR (B) of T cells treated with the indicated inhibitors or vehicle control during 48 hours of chronic TCR stimulation. Error bars represent SEM.

(C) Energy map visualization of OCR and ECAR in acutely stimulated T cells or chronically stimulated T cells treated with the indicated inhibitors or vehicle control during 48 hours of chronic TCR stimulation.

(D) Effect of indicated inhibitors on T cell proliferation during chronic TCR stimulation. Inhibitors: CRACi (p<0.0001), Calcineurini (p = 0.0027), PI3Ki (p<0.0001), p38i (p<0.0018), PKCi (p<0.0035), and mTORi (p<0.0001) after 144 hours for chronic stimulation.

(E–J) Population doublings of chronically stimulated T cells over time in the presence or absence of each inhibitor: CRACi (E), Calcineurini (F), PI3Ki (G), p38i (H), PKCi (I), and mTORi (J). Cells were treated at the specified concentrations during chronic TCR stimulation.

**Figure S3. MEKi-mediated suppression of terminal T cell exhaustion is dose- and time-dependent, but independent of nutrient restriction**

(A) Gene set enrichment analysis (GSEA) of calcineurin-regulated NFAT target genes in chronically stimulated T cells with or without MEK inhibition.

(B) Population doublings of acutely (left) or chronically (right) stimulated T cells treated with MEKi beginning 0, 48 or 96 hours following initial 48 hours of T cell activation.

(C) Representative flow cytometric plots of LAG-3 and TIM-3 expression in chronically stimulated T cells treated with MEKi at the indicated time points, highlighting % of terminally exhausted LAG-3^+^ TIM-3^+^ population.

(D) Fraction of m+2 malate derived from U-¹³C-glc in chronically stimulated T cells with or without MEKi treatment.

(E) Schematic highlighting non-canonical TCA cycle reactions.

(F) Ratio of m+2 malate to m+2 citrate in chronically stimulated T cells with or without MEKi treatment, reflecting canonical TCA cycle activity.

(G) Population doublings of acutely stimulated T cells cultured with or without MEKi in RPMI containing full (2 mM), or reduced (Low, 1 mM) glucose, or rescued (Low, 0.2 mM) glutamine.

(H) Population doublings of chronically stimulated T cells cultured with or without MEKi in RPMI containing full (2 mM) or reduced (Low, 0.2 mM) glutamine.

**Figure S4. MEK inhibition restricts chronic TCR stimulation-driven translation, ER stress responses, and terminal exhaustion in CD8^+^ T cells**

(A) Median cell volume (μm³) of acutely and chronically stimulated T cells treated with or without MEKi. Statistical significance determined using one-way ANOVA with multiple comparisons (p-values shown).

(B) Immunoblot analysis of ER stress response proteins ATF4 and BiP in acute and chronically stimulated T cells, with or without MEKi treatment. Vinculin serves as a loading control.

(C) Representative flow cytometry plots showing expression of exhaustion markers TIM-3 and Slamf6 on CD44^+^ PD-1^+^ Thy1.1^+^ P14 T cells isolated from LCMV Clone 13 infected mice treated with MEKi (Trametinib) or vehicle.

(D) Quantification of protein synthesis rates in PD-1^+^ CD44^+^ CD8^+^ T cells from LCMV Clone 13 infected mice treated with MEKi (Trametinib) or vehicle measured by O-propargyl-puromycin (OPP) mean fluorescence intensity (MFI) following 15 minutes pre-treatment with PBS or cycloheximide. Statistical significance was determined using two-way ANOVA with multiple comparisons (n=3 mice per group).

**Figure S5. Gating strategy for fluorescence activated cell sorting analysis.**

Gating was performed as shown: First, cells were gated by SSC-W versus FSC-W. Then, doublet exclusion was performed on cells gated by FSC-H versus FSC-A and SSC-H versus SSC-A. Viable cells were identified by FSC-A and Live/Dead GD510 exclusion. TCRb positivity was assessed by fluorescence in the BUV-805 channel. Finally, CD8 and CD4 positivity was assessed by fluorescence in the BUV-615 channel and BV-711 channel, respectively.

## STAR★METHODS

### KEY RESOURCES TABLE

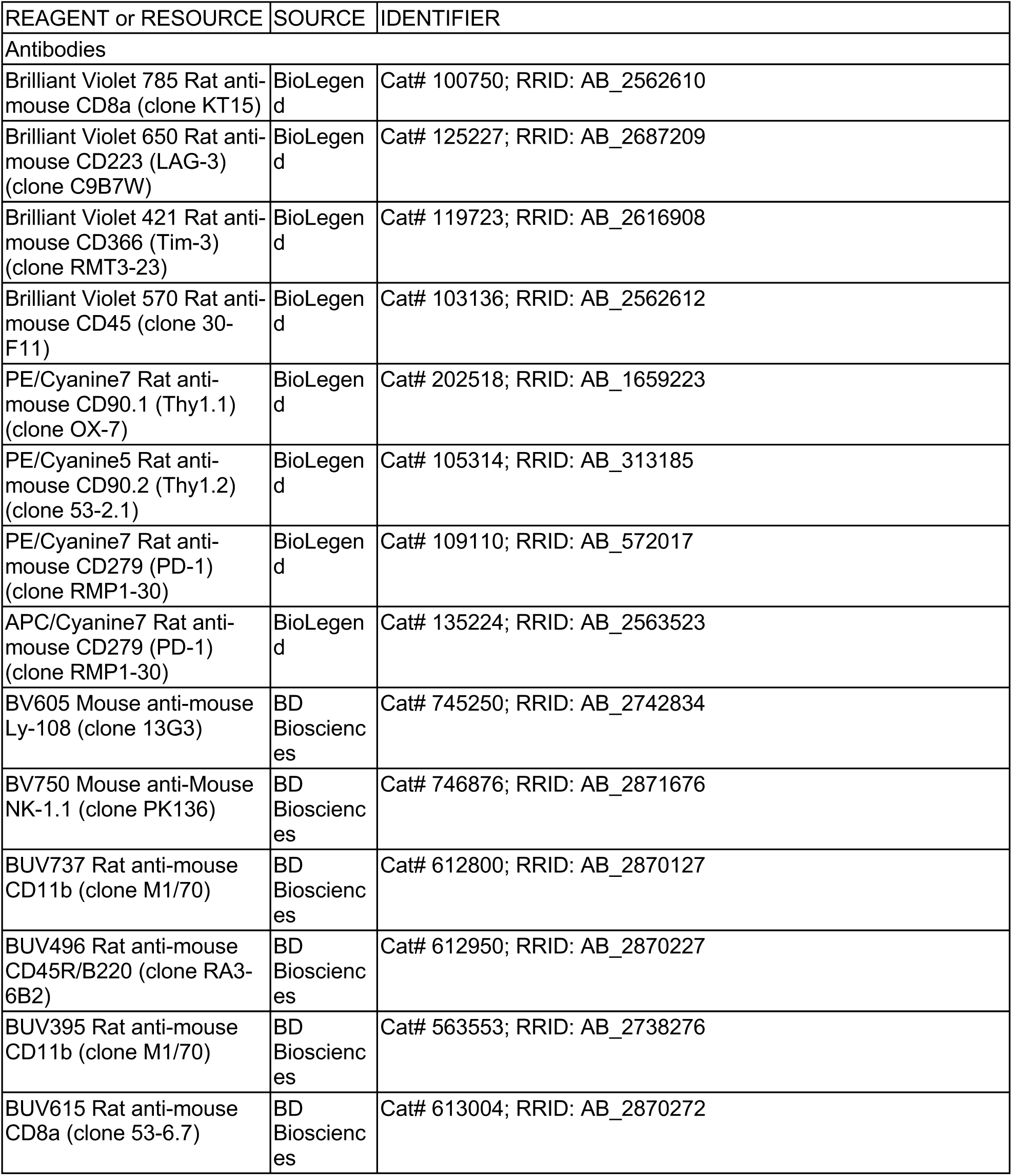
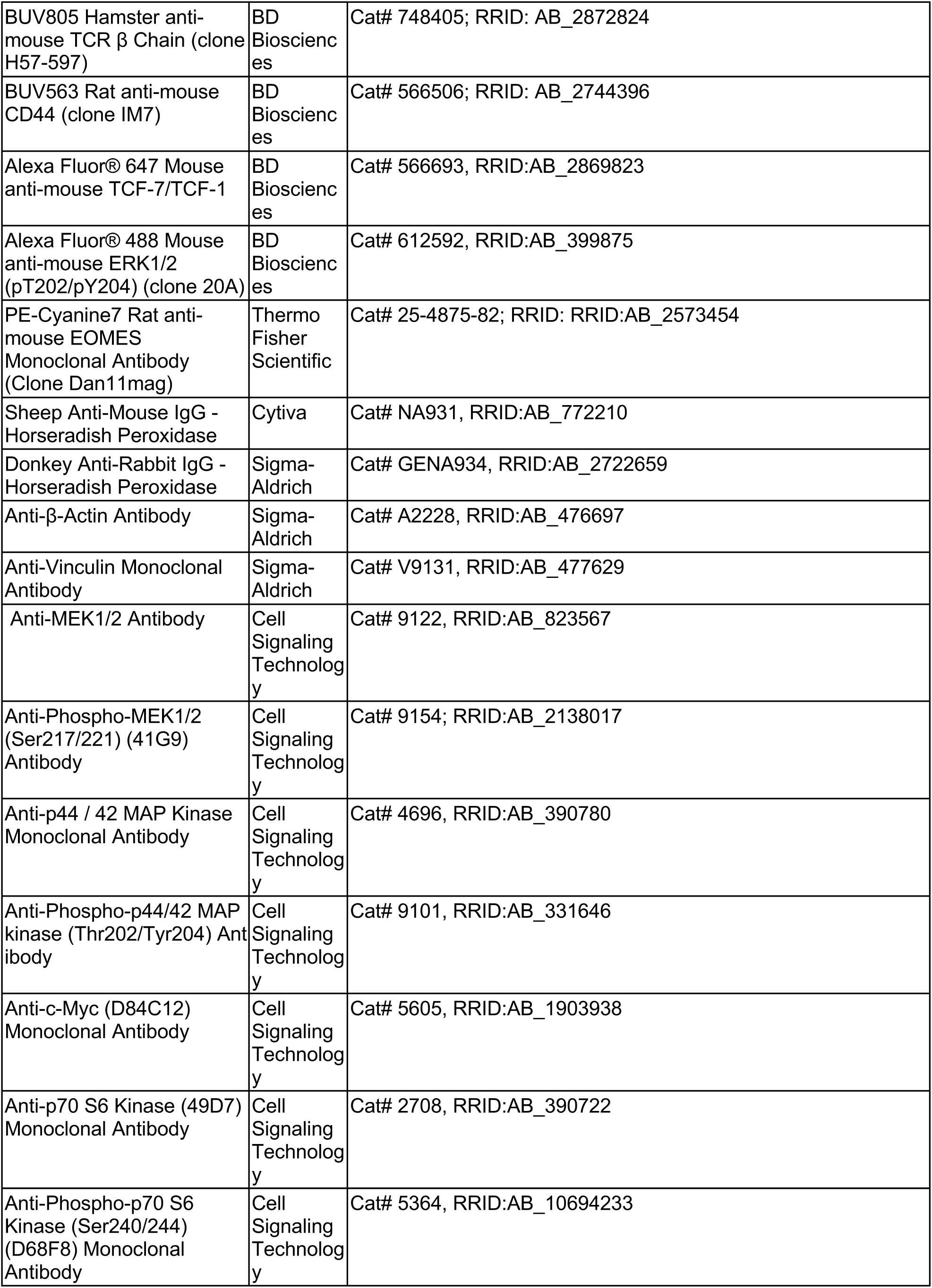
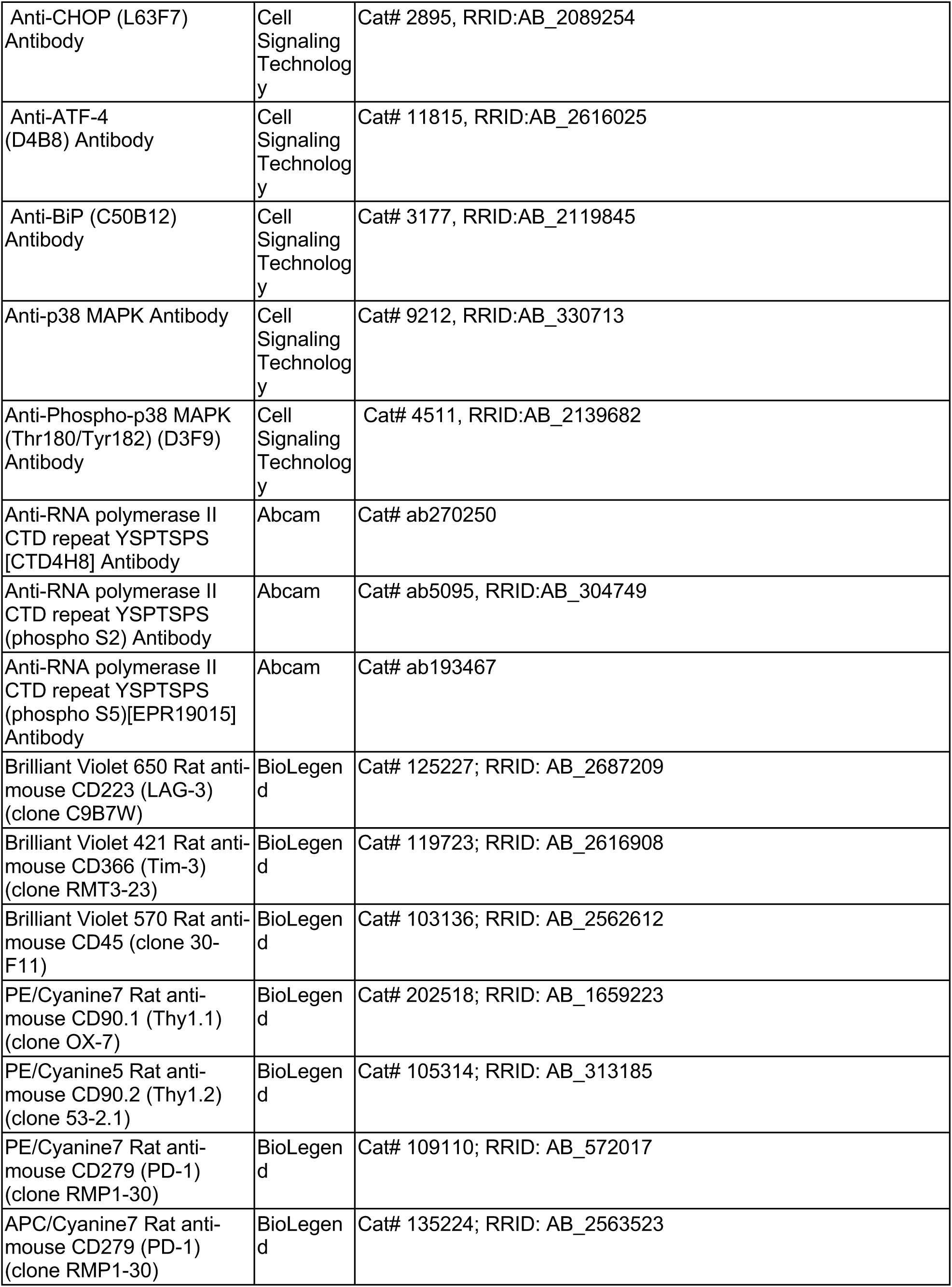
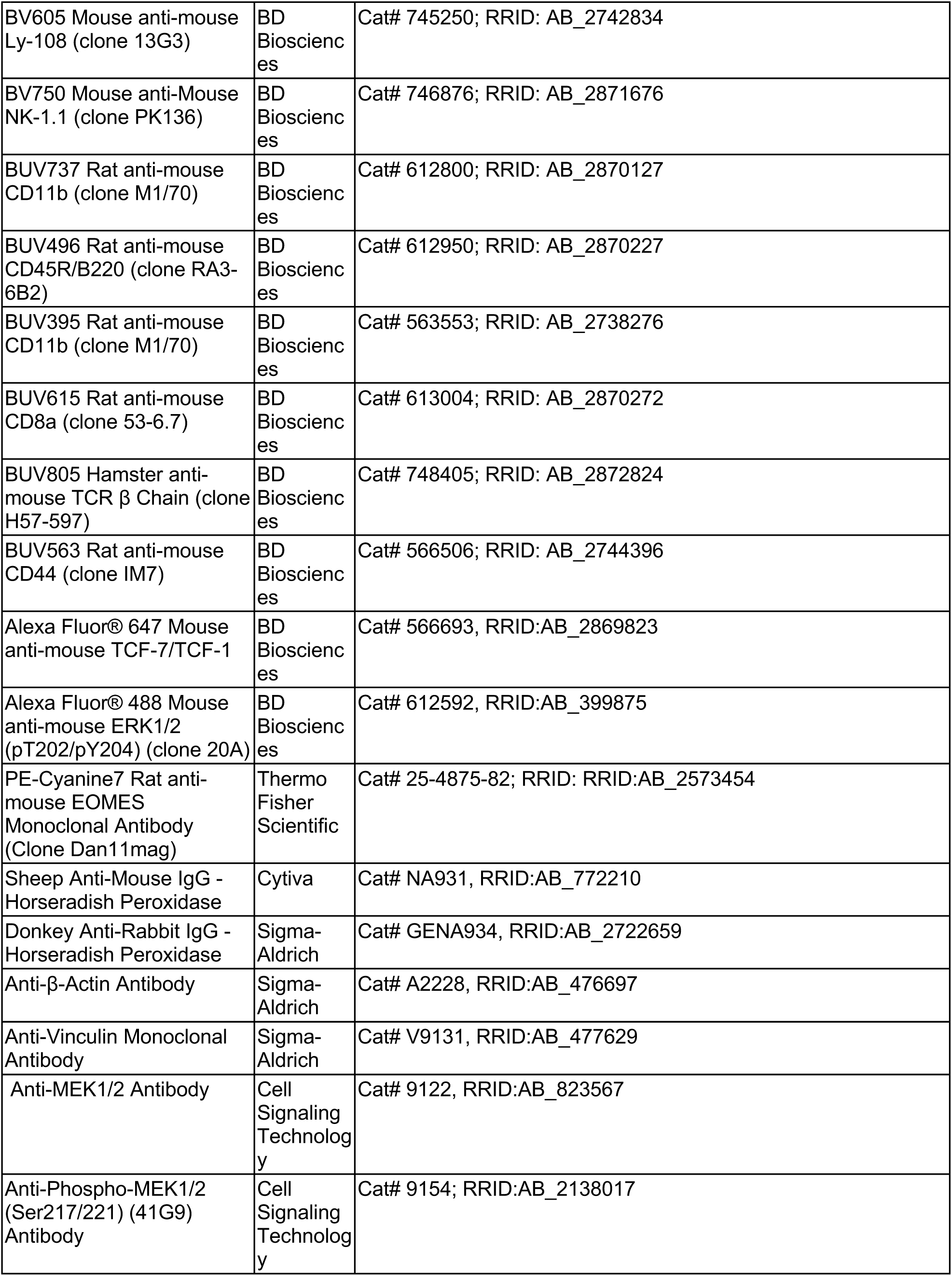
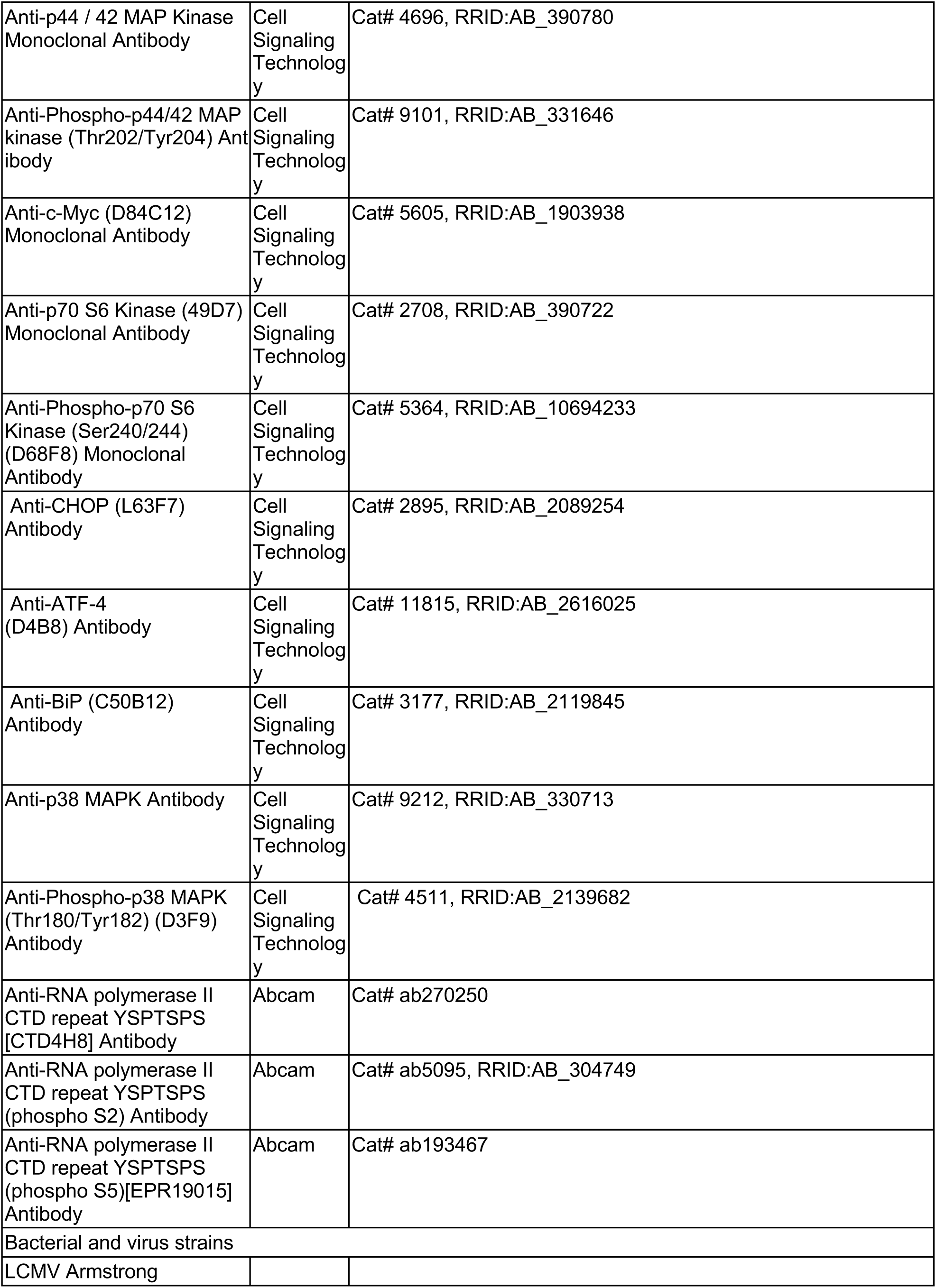
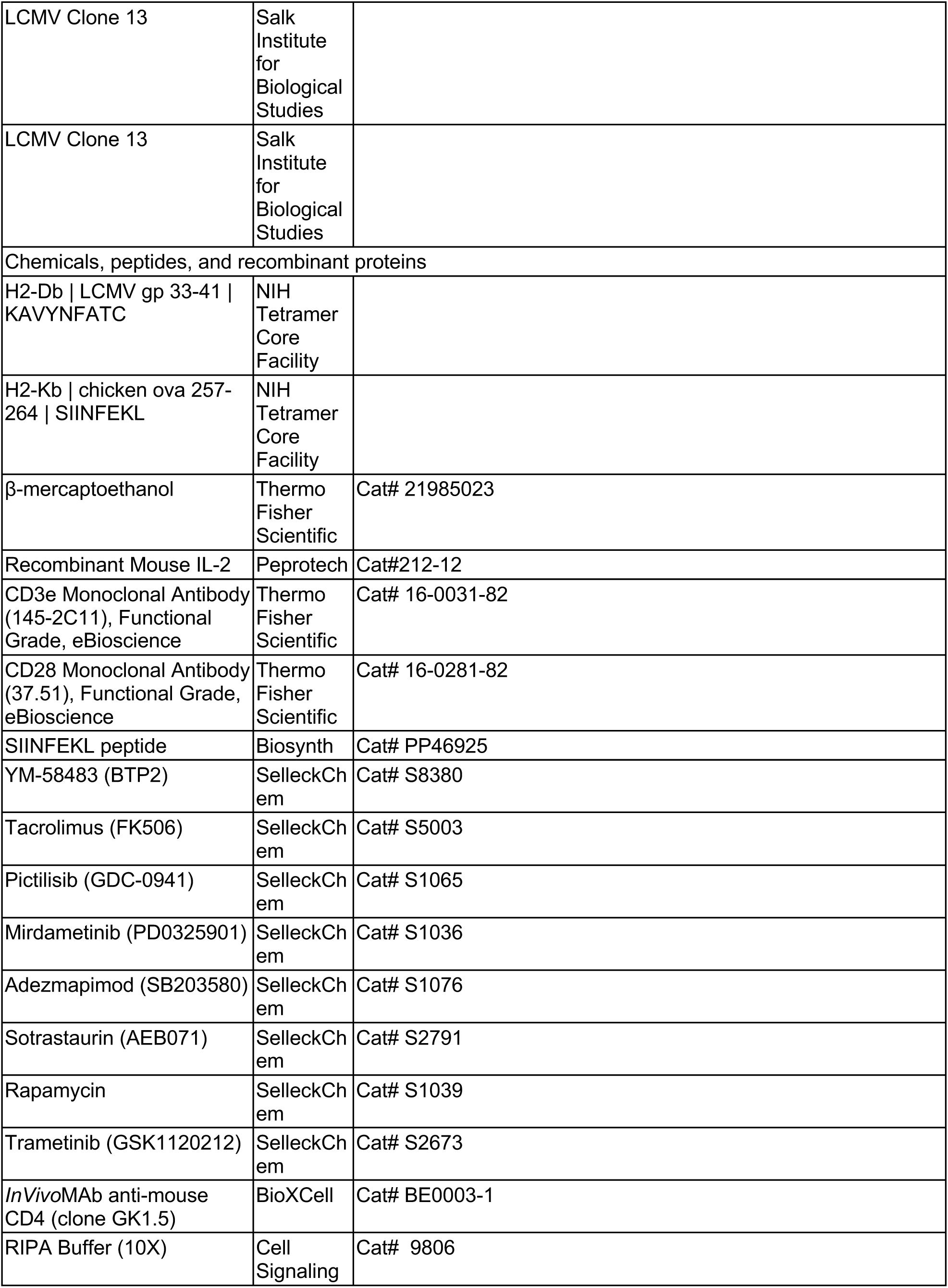
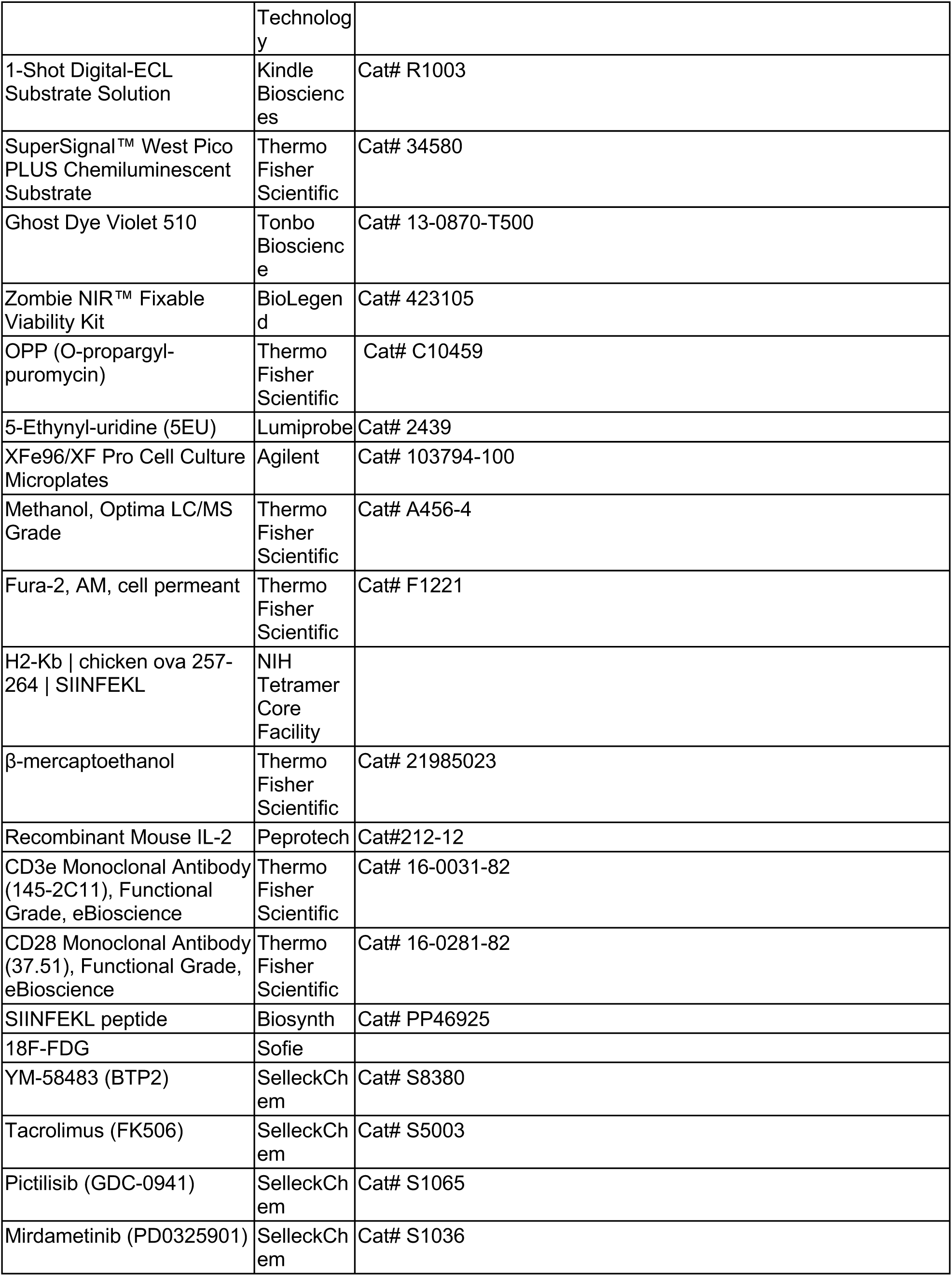
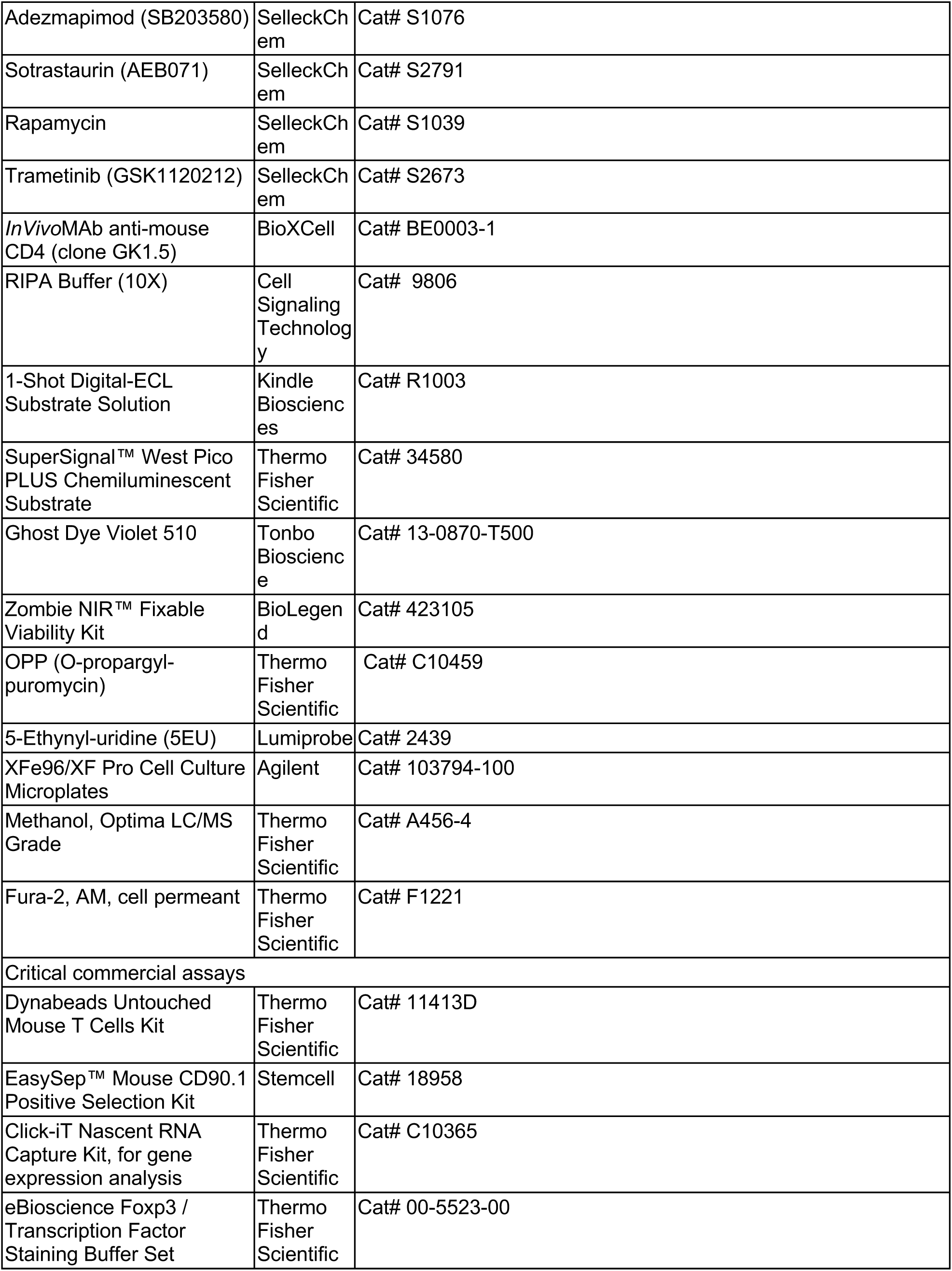
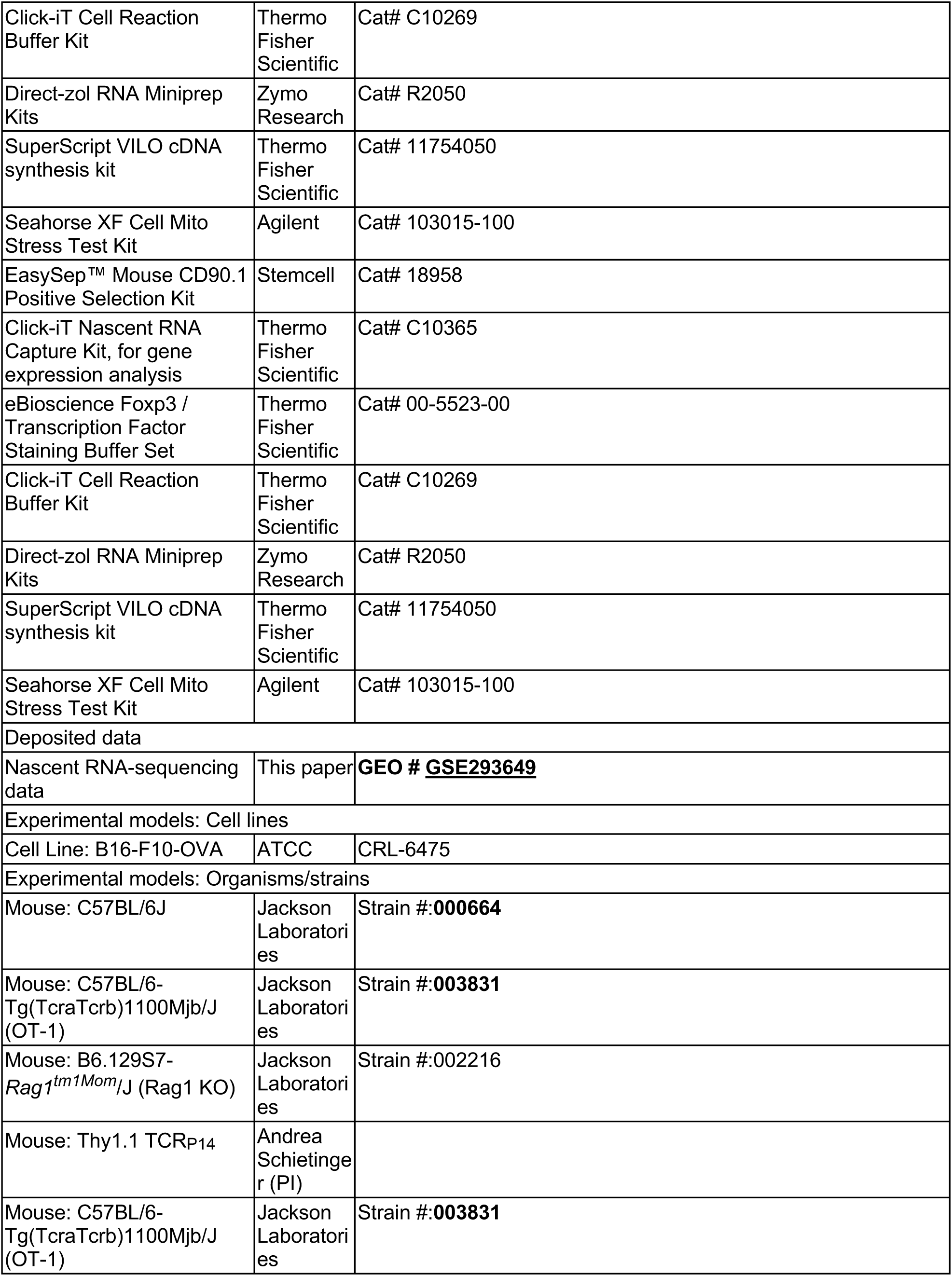
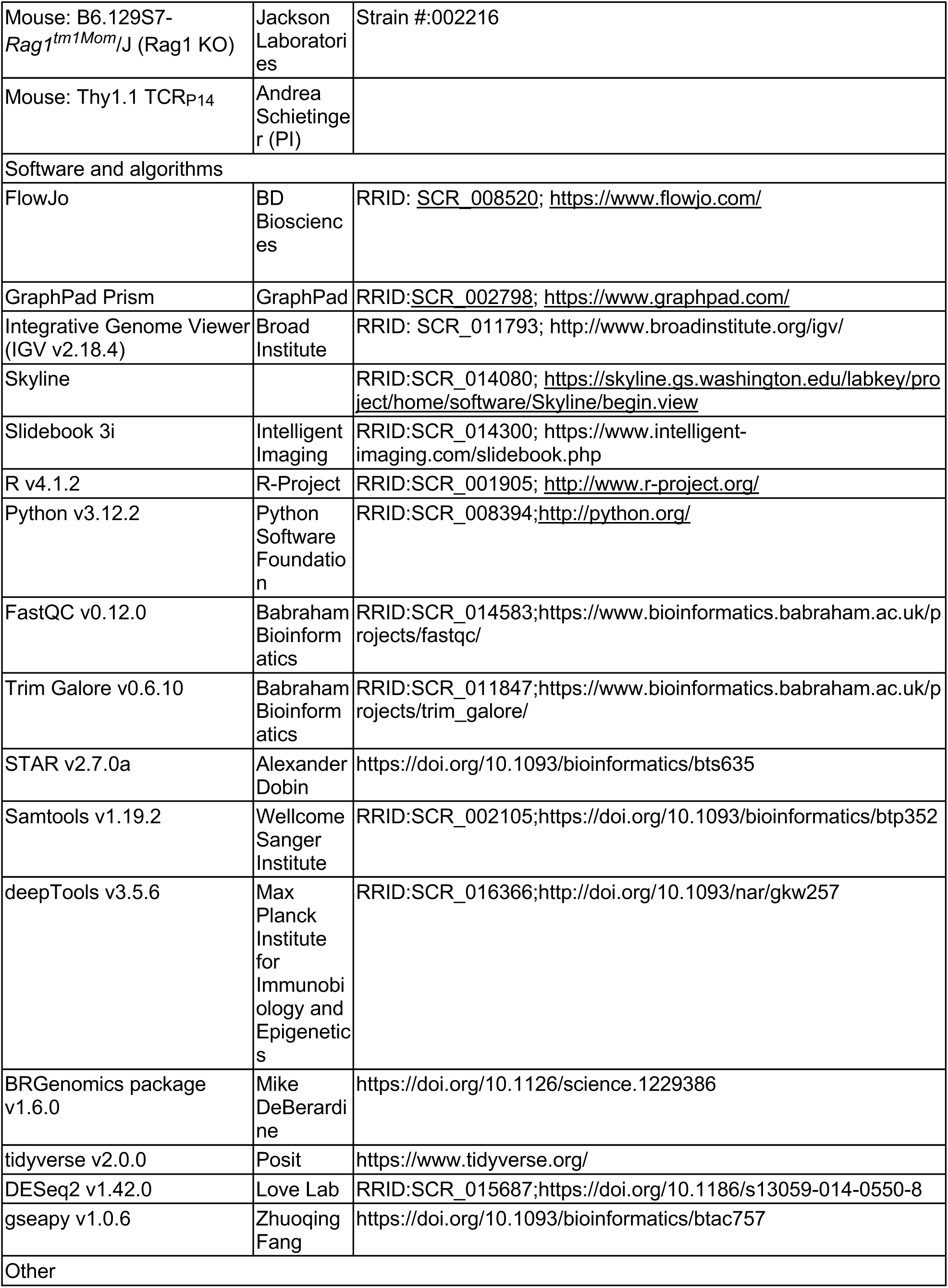
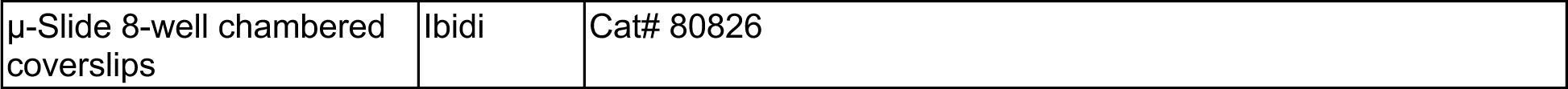

### EXPERIMENTAL MODEL AND STUDY PARTICIPANT DETAILS

#### Cell Lines

B16-OVA cell line was purchased from ATCC and cultured in DMEM supplemented with 10% heat-inactivated FBS. The cells were maintained at 37°C in 5% CO2, routinely tested for mycoplasma, and passaged no more than 15 times.

#### Mice

All mice were handled in accordance with NIH and American Association of Laboratory Animal Care standards. All mice used were on the C57BL/6 background. Mice were used at between 8 and 12 weeks of age unless otherwise indicated. Age and sex matched mice were used in each experiment. The following mice were housed and bred under specific pathogen-free conditions at Memorial Sloan Kettering Cancer Center (MSKCC) barrier facility: B6 (Strain # 000664), OT-1 (Strain # 003831), and Rag1 KO (Strain # 002216) mice were purchased from The Jackson Laboratory. P14 mice were kindly provided by Andrea Schietinger. All animal experiments were performed according to Memorial Sloan Kettering Cancer Center Institutional Animal Care and Use Committee (IACUC) guidelines (Protocol Number 20-10-012).

### METHOD DETAILS

#### T cell isolation, activation, and chronic stimulation

T cells were isolated from mouse spleens using Dynabeads (Thermo Fisher, #11413D) per the manufacturer’s instructions. Cells were cultured in RPMI-1640 with 10% fetal bovine serum (FBS), 2 mM L-glutamine, 50 μM β-mercaptoethanol (Thermo Fisher, #21985023), 50 U/mL Penicillin-Streptomycin, and 10 ng/mL murine IL-2 (Peprotech, #212-12), hereafter referred to as T cell media. For nutrient deprivation and isotope tracing, cells were cultured in glucose- or glutamine-free RPMI-1640 with 10% dialyzed FBS, 50 μM β-ME, 50 U/mL Penicillin-Streptomycin, 10 ng/mL murine IL-2, and specified glucose or glutamine concentrations. T cells were activated on plates coated with anti-CD3ε (3 μg/mL, Thermo Fisher, #16-0031-82) and anti-CD28 (1 μg/mL, #16-0281-82) for 48 hours, then maintained at 1 × 10^6 cells/mL in T cell media. Chronic stimulation was induced with plate-bound anti-CD3ε (3 μg/mL) for polyclonal CD8+ T cells or 100 nM SIINFEKL peptide (Biosynth, # PP46925) for OT-I transgenic CD8+ T cells, while acute conditions lacked these stimuli. T cells were cultured with fresh media changes every 48 hours and used for assays on day 4 (48 hours of chronic stimulation) or day 8 (144 hours of chronic stimulation).

#### *In vitro* drug treatments

T cells were treated with TCR pathway inhibitors post-activation, i.e., after 48 hours on anti-CD3ε- and anti-CD28-coated plates. The inhibitors used were BTP2 (CRAC channel inhibitor), Tacrolimus/FK506 (Calcineurin inhibitor), Pictilisib/GDC-0941 (PI3K inhibitor), Mirdametinib/PD0325901 (MEK inhibitor), SB203580 (p38 MAPK inhibitor), Sotrastaurin (PKC inhibitor), and Rapamycin (mTOR inhibitor), each tested at 1 μM, 100 nM, and 10 nM. Treatments were maintained with T cell media and inhibitor replacement every 48 hours. Inhibitor efficacy was confirmed by western blot analysis of downstream phosphorylation, and their effects on proliferation and metabolism were assessed.

#### Growth curve

Following activation, T cells were seeded in 24-well plates at 5 × 10⁵ cells/mL under different conditions. Every 48 hours, cells were collected and counted using a Beckman Coulter Counter with a volume gate of 75-4,000 femtoliters. After counting, 50% were re-plated in fresh medium in their respective conditions.

#### *In vivo* tumor treatments

Wildtype or Rag1 KO mice were subcutaneously injected with 2 × 10⁵ B16-OVA cells in PBS. When tumors reached ∼5 mm in diameter, 1 × 10⁶ OT-I T cells were adoptively transferred via retro-orbital injection. Mice were monitored daily for tumor growth. One-week post-transfer, they were treated with 1mg/kg Trametinib (SelleckChem) dissolved in 0.5% hydroxypropyl methylcellulose + 0.2% TWEEN 80, or vehicle, by oral gavage for five consecutive days. Mice were then euthanized, and tumors and lymph nodes were harvested for analysis.

#### Mouse LCMV infection

Lymphocytic Choriomeningitis Virus (LCMV) Armstrong (generated in-house) and Clone 13 stocks (Salk Research Institute) were quantified by plaque assay and stored as frozen aliquots. 1,000 Thy1.1+ P14 transgenic T cells were adoptively transferred into 6–7 weeks old wildtype mice via retro-orbital injection one day before infection. The virus was diluted in RPMI-1640 + 1% FBS, and either 2 × 10⁵ PFU of Armstrong were administered intraperitoneally, or 2 × 10⁶ PFU of LCMV Clone 13 intravenously. To deplete CD4+ T cells, 200 µg of anti-CD4 antibody (GK1.5; Bio X Cell) was injected intraperitoneally on days −1 and 1 of infection. Starting at day 12 or day 25 post-infection (depending on the experiment), mice were treated daily with 1 mg/kg Trametinib (SelleckChem) dissolved in 0.5% hydroxypropyl methyl cellulose + 0.2% TWEEN 80, or vehicle, by oral gavage for five consecutive days. Mice were then euthanized, and spleens and livers were harvested for analysis.

#### Measurement of glucose uptake *in vivo*

To assess glucose uptake by antigen-specific CD8+ T cells, LCMV-infected mice were fasted overnight and injected visa lateral tail vein with 18-fluorodeoxyglucose (^18^F-FDG). After 40 minutes, mice were euthanized, and spleens were harvested. Thy1.1^+^ P14 T cells were isolated using the EasySep Mouse CD90.1 Positive Selection Kit (StemCell Technologies, #18958) following the manufacturer’s protocol. The isolated cells were then assessed for ^18^F-FDG uptake by gamma counting using an automatic Wizard 2 γ-counter (Perkin Elmer). ^18^F background- and dead time-corrected count rates (count-per-minute (CPM)) values were obtained with an energy window of 400–600 keV. ^18^F count rates were converted to becquerels using a measured ^18^F system calibration factor decay-corrected to the time of injection, and reported as the percentage of injected dose (%ID). To account for differences in cell number, %ID was further normalized to the number of cells counted in each sample (%ID/cell). To improve the statistics, P14 T cells from three mice within the same treatment group were pooled for counting.

#### Western blot analysis

Protein lysates were extracted in RIPA buffer (Cell Signaling Technology, #9806), separated by SDS-PAGE, and transferred to nitrocellulose membranes (Bio-Rad). Membranes were blocked with 5% milk in Tris-buffered saline with 0.1% Tween-20 (TBST) and incubated overnight at 4°C with primary antibodies diluted in 5% bovine serum albumin (BSA) in TBST. After three TBST washes, membranes were incubated with horseradish peroxidase (HRP)-conjugated secondary antibodies (GE Healthcare, anti-mouse IgG-HRP, GENA931; anti-rabbit IgG-HRP, GENA934; 1:5,000) diluted in 5% milk in TBST for 1 hour at room temperature. Enhanced chemiluminescence (ECL) substrate (1-Shot Digital-ECL, Kindle Biosciences #R1003 or SuperSignal West Pico PLUS Chemiluminescent Substrate, Thermo Fisher #34580) was applied, and membranes were imaged using the ChemiDoc Touch Imaging System (Bio-Rad). Primary antibodies were used at 1:1,000 unless noted otherwise: MEK1/2 (CST #9122), p-MEK1/2 (Ser217/221) (CST #9154), ERK1/2 (CST #4696), p-ERK1/2 (Thr202/Tyr204) (CST #9101), c-Myc (CST #5605), S6 Ribosomal Protein (CST #2708), p-S6 Ribosomal Protein (Ser240/244) (CST #9234), CHOP (CST #2895), ATF-4 (CST #11815), BiP (CST #3177), p-p38 MAPK (Thr180/Tyr182) (CST #4511), p38 MAPK (CST #9228), RNA Polymerase II CTD repeat YSPTSPS (Abcam ab270250), CTD repeat p-Ser2 (Abcam ab5095), CTD repeat p-Ser5 (Abcam ab193467), Vinculin (Sigma #V9131), and β-Actin (Sigma #A2228, used at 1:20,000).

#### Immunophenotyping

Cells were labeled with a viability dye (Ghost Dye 510, Tonbo #13-0870-T500 or Zombie NIR, BioLegend #423105) for 10 minutes at 4°C, then stained with surface markers for 20 minutes at room temperature. For intracellular staining, cells were washed and fixed for 30 minutes at room temperature using the FoxP3 Fixation/Permeabilization Kit (Thermo Fisher, #00-5523-00). Intracellular markers were then stained overnight at 4°C, followed by fixation with 4% formaldehyde in PBS. Samples were recorded on a Cytek Aurora, and data were processed using FlowJo v10.10. Antibodies used are listed in the Key Resources Table, with gating strategies shown in Figure S5.

#### Measurement of nascent protein synthesis

For cell culture assays, 5 × 10⁵ T cells were seeded in 500 μL of T cell media per well in a 12-well plate. Cells were incubated with 10 μM O-Propargyl-Puromycin (OP-Puro; Thermo Fisher C10459) for 30 minutes at 37°C to label nascent proteins. Negative controls were pretreated with 10 μg/mL Cycloheximide. For in vivo experiments, mice were injected intraperitoneally with OP-Puro at 49.5 mg/kg, and the negative control group was pretreated with 30 mg/kg Cycloheximide intraperitoneally, 10 minutes prior to OP-Puro injection^63^. After the designated incorporation period, tissues were harvested, and single-cell suspensions were prepared.

Labeled cells (from both *in vitro* and *in vivo* experiments) were stained for viability and surface markers, then fixed and permeabilized using the Click-iT™ Protein Reaction protocol (Thermo Fisher, #C10269). OP-Puro incorporation was detected with Alexa Fluor 647 dye via Click-iT reaction, followed by flow cytometry analysis on a Cytek Aurora. Mean fluorescence intensity was quantified using FlowJo v10.10.

#### Nascent RNA quantification and sequencing

Nascent RNA was labeled in live cells and isolated using the Click-iT™ Nascent RNA Capture Kit (Thermo Fisher, Cat. No. C10365) following the manufacturer’s protocol. Briefly, 1 × 10⁶ T-cells were seeded in 500 µL of T-cell media per well in a 12-well plate. Actinomycin D (100 pM) was added to the negative control wells for 12 hours prior to labeling. Cells were then incubated with 0.2 mM 5-Ethynyl uridine (5-EU) (Lumiprobe, #2439) at 37°C for 30 minutes to label newly synthesized RNA. After harvesting, cells were centrifuged at 250g for 5 minutes, and total RNA was isolated using the Direct-zol RNA Miniprep Kit (Zymo Research, #R2050).

For sequencing nascent RNA, the 5-EU-labeled nascent RNA was biotinylated through a click reaction with biotin-azide for 1 hour at room temperature, bound to Dynabeads MyOne Streptavidin T1 magnetic beads, and isolated for cDNA synthesis. Reverse transcription was performed using the SuperScript VILO kit (Thermo Fisher, Cat. No. 11754050). The resulting cDNA was purified and sequenced. Sequencing was performed by the MSK Integrated Genomics Operation Core. For quantification in a limited number of cells, 5-EU-labeled nascent RNA was clicked to Alexa 488-azide and measured by flow cytometry.

#### Bulk RNA-seq

RNA was isolated from T cells using the Direct-zol RNA Miniprep Kit (Zymo Research, #R2050) following the manufacturer’s instructions. RNA quantity and integrity were assessed using a NanoDrop spectrophotometer and Agilent Bioanalyzer, respectively, before sequencing. Library preparation and sequencing were performed by the MSK Integrated Genomics Operation Core.

#### Extracellular flux analysis

Extracellular acidification rate (ECAR) and oxygen consumption rate (OCR) were measured using the Seahorse XFe96 Extracellular Flux Analyzer (Agilent Technologies) following established protocols described in Vardhana et al^19^. T cells were plated in poly-L-Lysine-coated XF 96-well plates (Agilent, #103794-100) at 2 × 10⁵ cells per well in assay medium (phenol red-free RPMI containing 10 mM glucose, 2 mM L-glutamine, and 1 mM sodium pyruvate). Baseline ECAR and OCR were recorded, followed by sequential injections of 1 μM oligomycin, 1 μM FCCP, and 0.5 μM rotenone/antimycin mix. In some experiments, 2 μM UK5099 or 3 μM BPTES, or 10 μg/mL Cycloheximide, were injected at the start to assess substrate oxidation dependency. Mitochondrial ATP production was calculated by subtracting the respiration rate after oligomycin injection from the baseline rate, while ATP production from glycolysis was inferred from the Proton Efflux Rate (PER) measurements.

#### Measurement of nutrient uptake and excretion

Cells were plated in 12-well plates at a density of 1 × 10⁶ cells per mL of T cell media per well, with additional wells containing media alone as references for original metabolite concentrations. After 24 hours, the media was collected and centrifuged at 300g for 3 minutes 4°C to remove cells and debris. The concentrations of glucose, glutamine, and lactate in the media were analyzed using a 2950D Biochemistry Analyzer (YSI Life Sciences), as described previously^19^. The metabolite concentrations were normalized to cell number, medium volume, and incubation time.

#### Stable isotope tracing

T cells were washed with PBS and seeded at a concentration of 2 × 10⁶ cells/mL in glucose-deficient RPMI, supplemented with 10% dialyzed FBS, 2 mM L-glutamine, 50 μM β-mercaptoethanol, 50 U/mL Penicillin-Streptomycin, and 10 ng/mL murine IL-2, along with either 2 g/L of uniformly labeled ¹³C- or ¹²C-glucose. At the designated time points, cells were harvested and metabolites were extracted with 80:20 methanol:water, then stored at −80°C overnight. After incubation, lysates were centrifuged at 20,000g for 20 minutes at 4°C to remove proteins. Supernatants were collected, dried down using a Genevac evaporator, and stored at −80°C until the time of analysis. For isotope-labeling analysis, liquid chromatography-mass spectrometry (LC-MS) methods were employed as described previously^19^. Briefly, dried extracts were resuspended in 100 µL of water for ion pair (IP) liquid chromatography separation and in 60 µL of 60:40 acetonitrile:water for hydrophilic interaction liquid chromatography (HILIC).

For TCA metabolites and nucleotide phosphates, IP separation of analytes was accomplished with a XSelect HSS T3 column (150 × 2.1 mm, 3.5 µm particle size) (Waters Corp.), paired to a 6230 TOF MS (Agilent Technologies) via negative mode ionization. A reverse-phase gradient of solvent A (5 mM octylamine and 5 mM acetic acid in water) and solvent B (5 mM octylamine and 5 mM acetic acid in 90:10 methanol:water) was applied with the following LC timing: 0–3.5 min, 1% B; 4–15 min, 35% B; 20–22 min, 100% B; and 22–27 min, 1% B, infused with a post-column flow consisting of a blend of acetone:DMSO (90:10) at 300 µL/min.

For polar analytes, a Waters Acquity UPLC BEH Amide column (150 × 2.1 mm, 1.7 µm particle size) was used for HILIC LC separation. The gradient used was: 0–9 min, 95% B; 9–13 min, 60% B; 13–14 min, 30% B; and 14.5–20 min, 95% B, where solvent A was composed of 10 mM NH4 acetate, 10% acetonitrile, and 0.2% acetic acid, and solvent B was 10 mM NH4 acetate, 90% acetonitrile, and 0.2% acetic acid. MS analysis was performed on a 6545 Q-TOF MS (Agilent Technologies) in positive ionization mode.

#### Steady-state metabolite profiling

T cells were seeded at a concentration of 2 × 10⁶ cells/mL in T cell media. After 2 days in culture, cells were harvested and metabolites were extracted with 80:20 methanol:water, then stored at −80°C overnight. After incubation, lysates were centrifuged at 20,000g for 20 minutes at 4°C to remove proteins. Supernatants were collected, dried down using a Genevac evaporator, and stored at −80°C until the time of analysis. For steady-state metabolomics, these dried extracts were resuspended in 50 μL of mobile phase A and incubated on wet ice for 20 minutes, being vortexed every 5 minutes. Samples were centrifuged at 20,000 x g for 20 minutes at 4°C, and 40 µL of the supernatant was transferred into LC vials for injection.

Ion pair LC-MS/MS analysis was performed using an Agilent 1290 Infinity II LC system coupled to an Agilent 6470 Triple Quadrupole (Agilent Technologies). Chromatographic separation was achieved using a Zorbax RRHD Extend-C18 column (150 mm × 2.1 mm, 1.8 μm, Agilent Technologies), and using a gradient of solvent A (10 mM tributylamine and 15 mM acetic acid in 97:3 water:methanol) and solvent B (10 mM tributylamine and 15 mM acetic acid in methanol) according to the manufacturer’s instructions (MassHunter Metabolomics dMRM Database and Method, Agilent Technologies).

#### Calcium flux measurements

µ-Slide 8-well chambered coverslips (Ibidi, #80826) were coated with biotin-poly-L-lysine, blocked with 1% BSA in PBS, and incubated with 100 µg/mL streptavidin in 1% BSA for 1 hour. The wells were then incubated with anti-CD3e (clone 145-2C11) and ICAM-1 for 1 hour. Fura-2 dye (Thermo Fisher, #F1221) was added to 1 × 10⁶ T cells/mL at a concentration of 4 µM and incubated at 37°C for 30 minutes. The cells were washed twice by centrifugation (500g for 2 minutes) and resuspension in dye-free, phenol red-free T cell media. Images were acquired on an inverted fluorescence microscope (Olympus IX-81) fitted with a 20× objective lens (Olympus) using the Slidebook software (3i). Fura-2 imaging was performed by capturing images of 510-nm emission at 340- and 380-nm excitation every 30 s for 30 min at multiple positions per well. Population Ca^2+^ responses were analyzed on Slidebook by applying image background correction, generating cell masks using Otsu thresholding and size exclusion, and measuring cross-channel ratiometric intensity values across the segment masks in each video.

### QUANTIFICATION AND STATISTICAL ANALYSIS

#### Nascent RNA-seq Analysis

Nascent RNA-seq samples were sequenced using the Illumina Novaseq 6000 platform, generating 100-base pair paired-end reads. FASTQ files were quality-checked using FastQC v0.12.0 and trimmed with Trim Galore v0.6.10 to remove adapter sequences and read ends with Phred scores below 15. STAR v2.7.0a was used to align the FASTQ files to the mm10 reference genome. The resulting SAM files were sorted using the sort function from Samtools v1.19.2 and indexed using the index function to generate BAM index files. Stranded bigWig files were generated using the bamCoverage function from deepTools v3.5.6. The BRGenomics package v1.6.0 was then used in R v4.1.2 to construct gene body and pause region count matrices, which were subsequently used to create pausing index tables with tidyverse v2.0.0. Downstream differential expression analysis was performed using DESeq2 v1.42.0. Gene set enrichment analysis was conducted using the prerank module from gseapy v1.0.6.

#### Metabolomic Analysis

Data analysis and relative quantification were performed with Skyline software (21).

#### Statistical analysis

Graphs and statistical analyses were generated and performed using GraphPad Prism Version 10.2.2 (341). Error bars, P values and statistical tests are reported in figure legends.

