## Supplementary figures and images for "MEK-dependent bioenergetic demand drives terminal CD8^+^ T cell exhaustion"

### Supplementary Figures 1-5

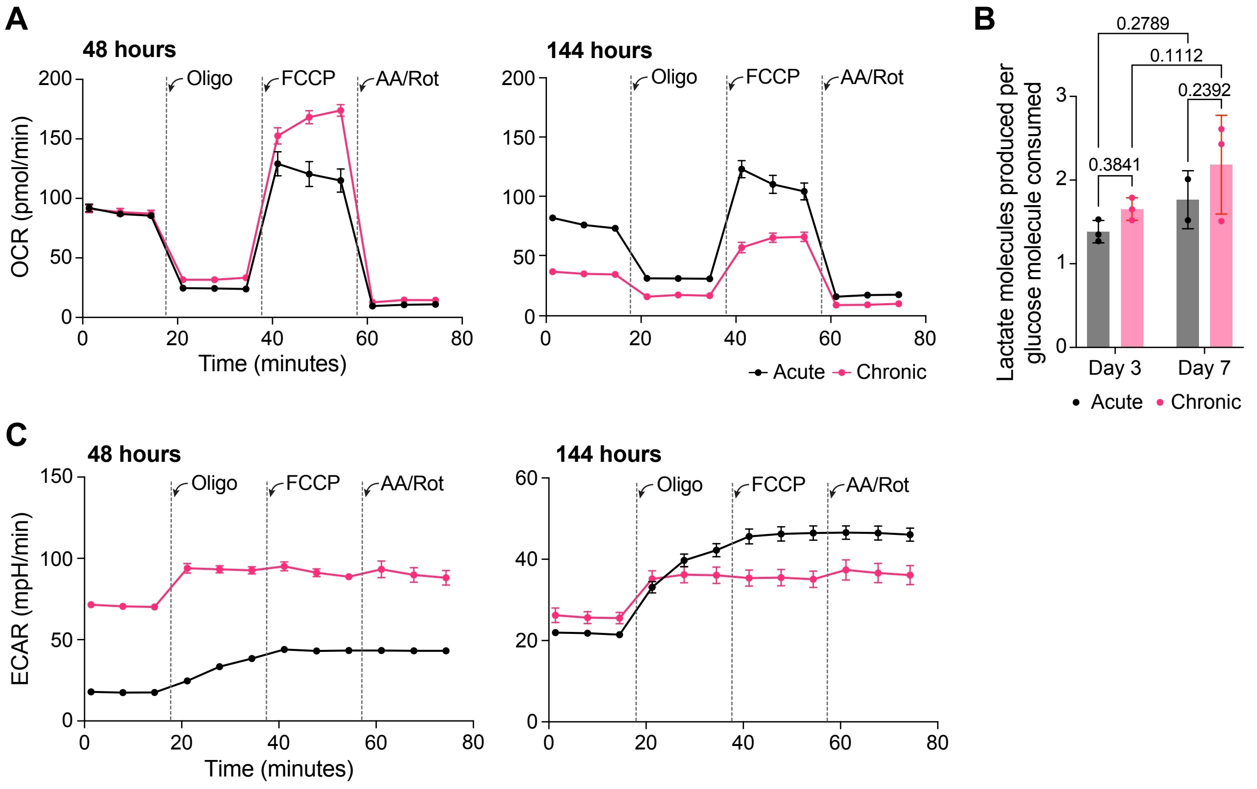

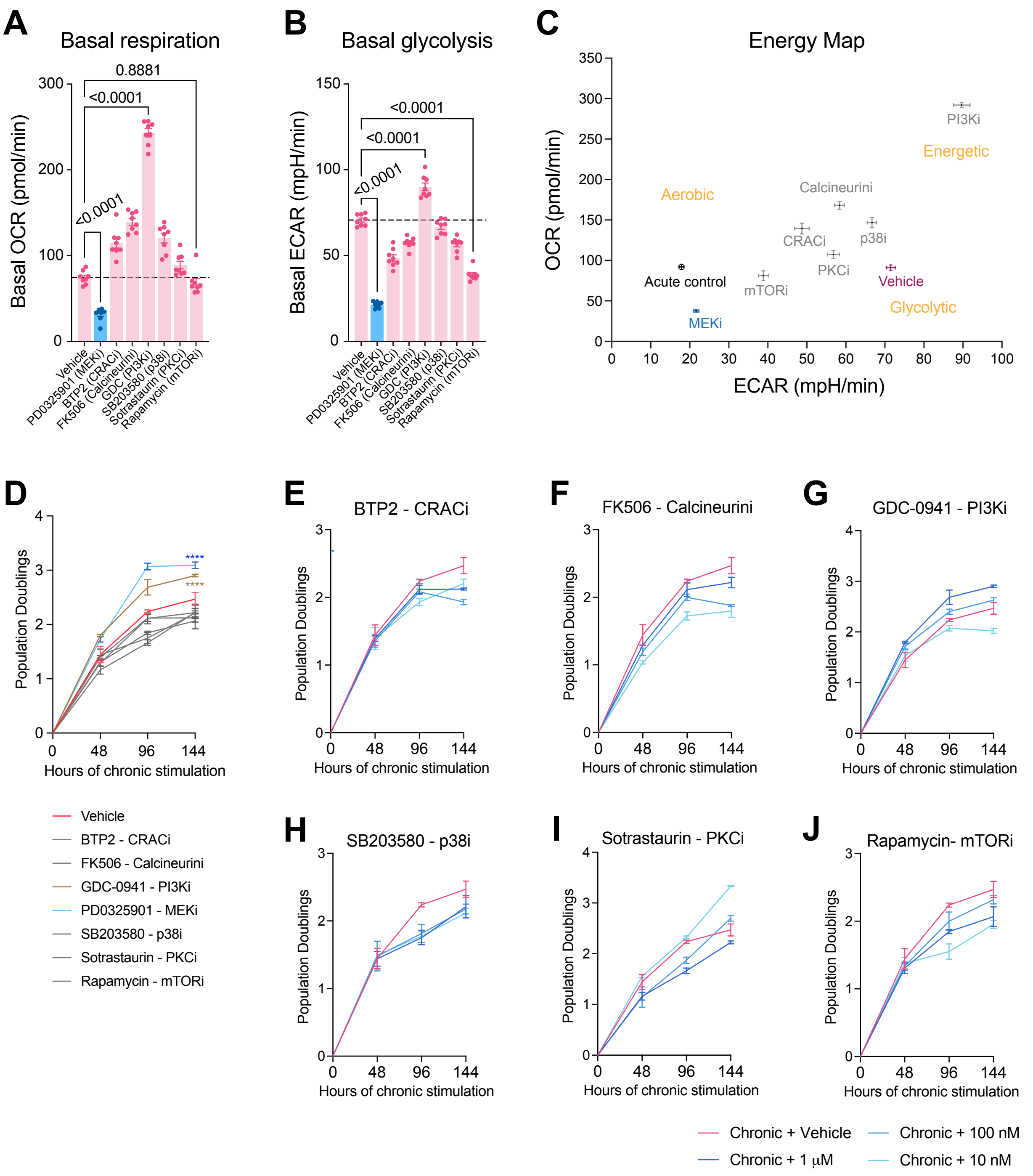

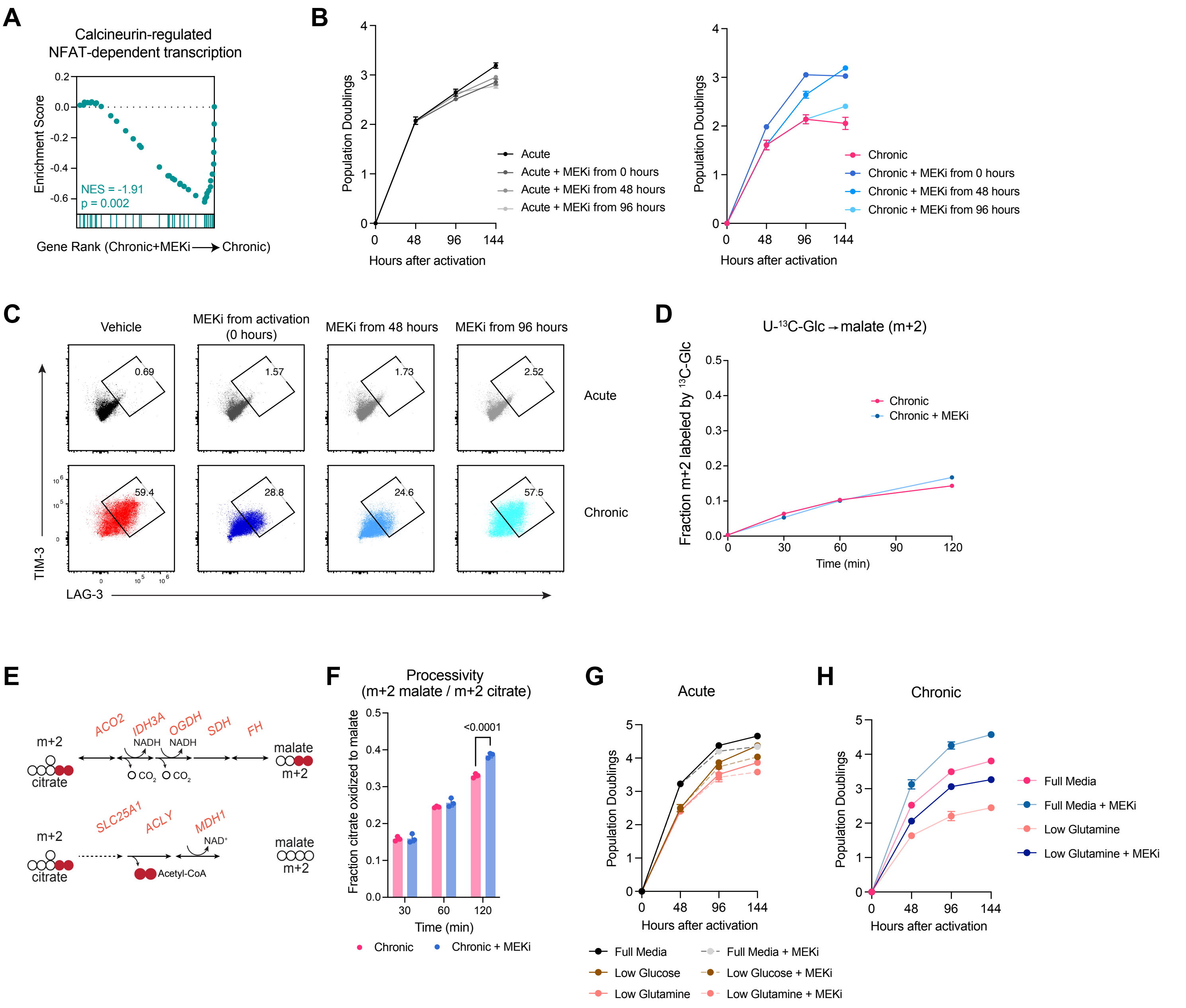

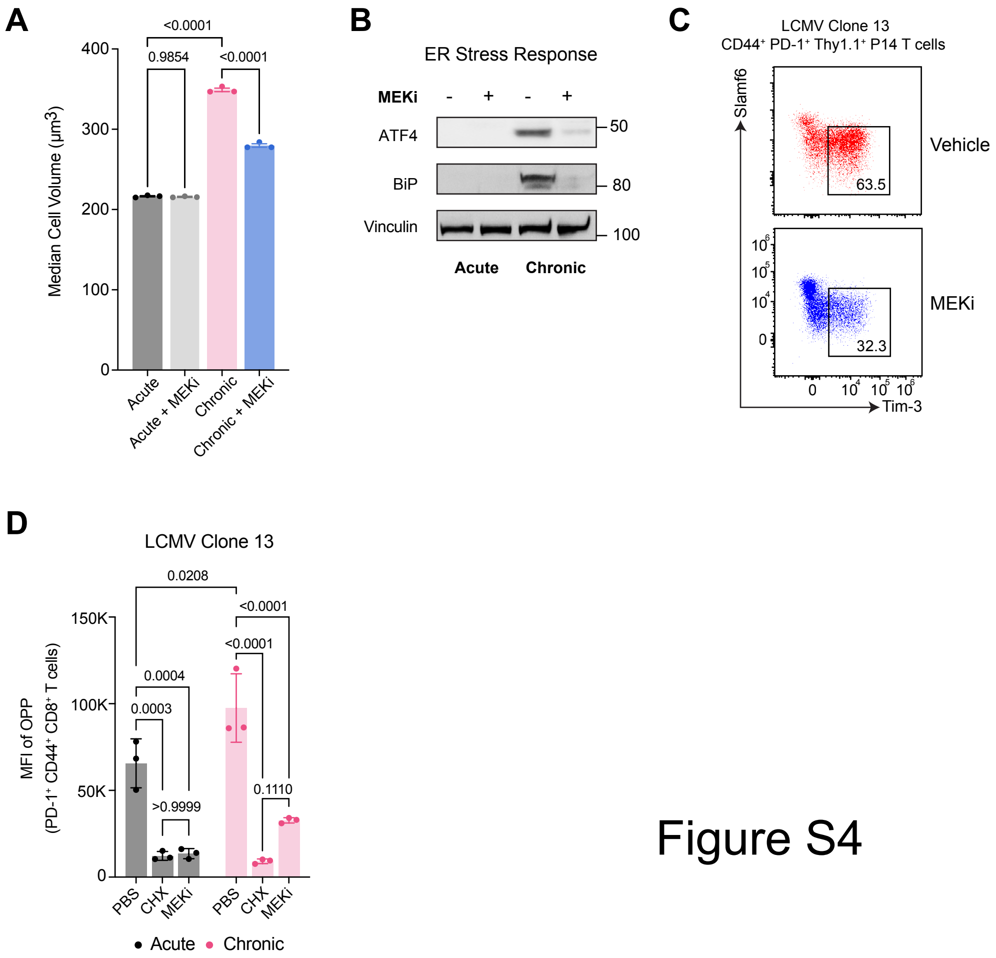

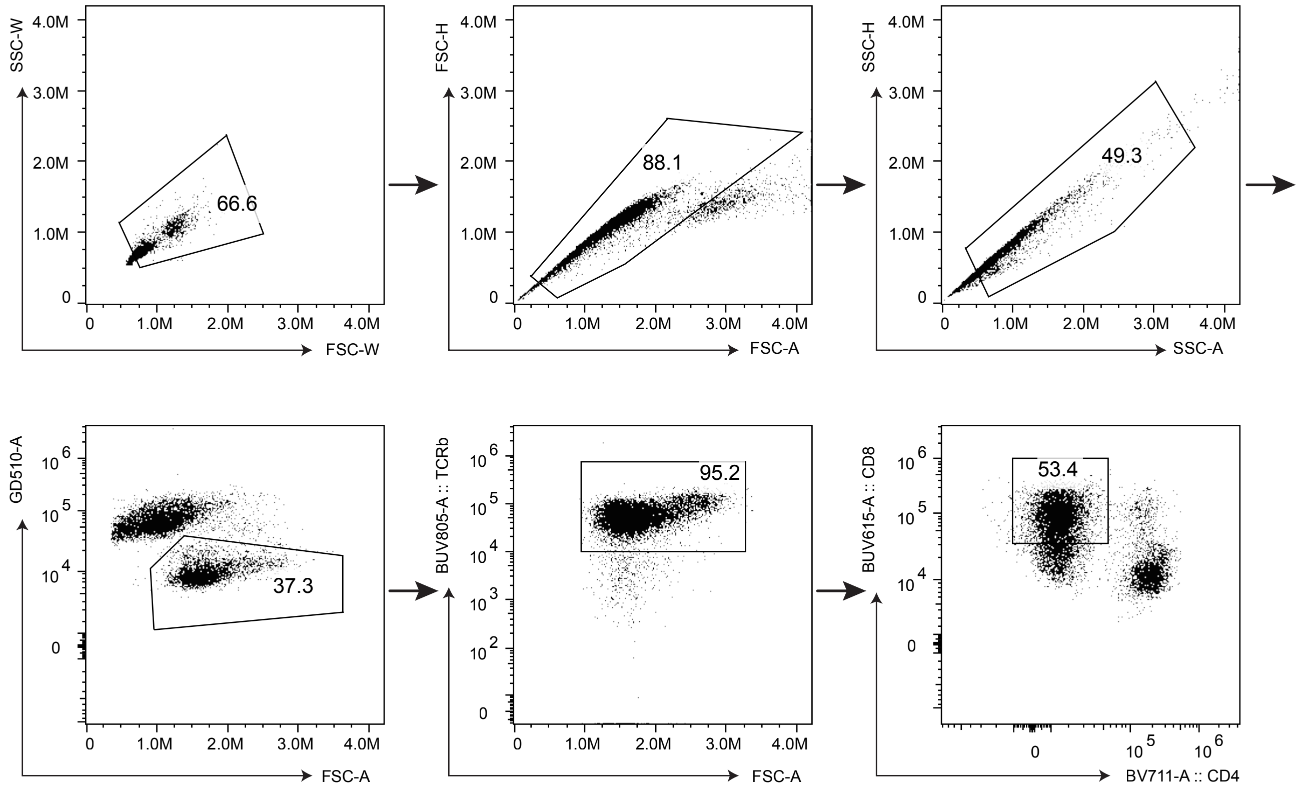
